# A new neural code for position: generalizable spatial maps in mouse medial frontal cortex

**DOI:** 10.64898/2026.10.21.759829

**Authors:** Rajyashree Sen, Lorenzo Posani, Alexander Medoff, Baihe Zhang, LF Abbott, Richard Axel, Stefano Fusi

## Abstract

Navigation requires the brain to encode spatial relationships that support extrapolation from explored to unexplored regions of space. We identified a novel neural code for space in the mouse medial frontal cortex (mFC) that is well suited for this form of spatial generalization. Using calcium imaging of excitatory neurons in mFC, hippocampus (dorsal CA1), and lateroventral orbitofrontal cortex (lvOFC) as mice explored a square arena with landmarks, we found that mFC activity supports extrapolation. Linear regression of the animal’s *x* or *y* coordinates trained in one region of the arena generalized to a completely different region. This property was observed in mFC but not in dCA1 or lvOFC. A model in which neurons encode the squared distance from the animal to multiple landmarks explains several properties of the geometry of the mFC representation, including extrapolation. The neural map that we observed in the frontal cortex may provide a basis for vector computation in navigation and complex route planning.

## Introduction

Animals navigate by building internal models that allow them to infer relationships between locations, estimate distances, and plan routes through regions they have never explored. Neural representations of spatial location have been studied intensively in two populations, place cells in the hippocampus (1, 2) and grid cells in the entorhinal cortex (3), but have now been described in many other brain areas (4–7). In its simplest form, an internal model of space that provides a basis for navigation and complex planning resembles a map in an atlas, and preserves the geometry of physical space. Such a map is compositional: spatial displacements can be combined as building blocks to construct routes through space. This is possible because, in a map, a given displacement represents the same change in position, regardless of where the movement begins.

Consider the street grid of Manhattan in New York City (Fig. 1a). Moving 8 blocks uptown along 8th Avenue takes us from 51st Street to Central Park at 59th Street. This movement defines a displacement of 8 blocks northward, represented by a vector on the map. The same displacement can be applied along a different avenue: moving 8 blocks uptown along 5th Avenue would also take us from 51st Street to 59th Street, even if we had never explored that avenue. This ability to apply a displacement independently of its starting location arises because a map captures the underlying geometry of space rather than representing a collection of individual locations or routes.

**Fig. 1.**
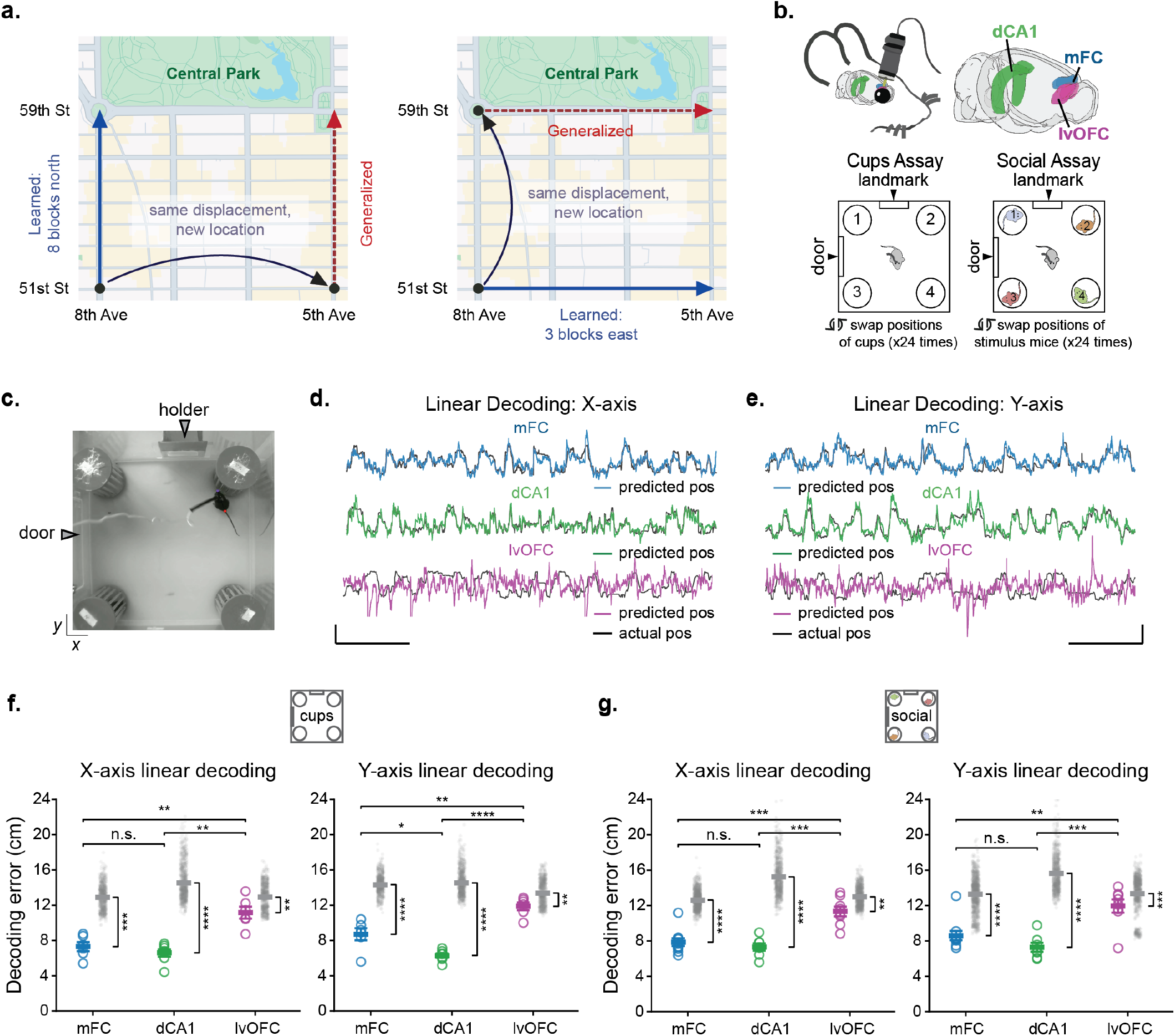
A framework and experimental paradigm for studying neural maps of space. **a**, Schematic illustrating spatial generalization using the Manhattan street grid. A displacement learned along one avenue, such as moving eight blocks uptown from 51st Street to 59th Street, can be applied at a new location along another avenue. Likewise, an east-west displacement learned between 8th and 5th Avenues at 51st Street can be generalized to the same displacement at 59th Street. **b**, Top, mice were injected with an AAV expressing GCaMP6f under the CaMKII*α* promoter and implanted with a GRIN lens over one of three brain regions - the medial frontal cortex (mFC), dorsal CA1 of the hippocampus (dCA1), or lateroventral orbitofrontal cortex (lvOFC) - for one-photon calcium imaging of excitatory neurons. Bottom, mice explored an arena containing four salient corner landmarks: either four identical cups positioned in the corners (cups assay) or four individually housed stimulus mice (social assay). In both assays, the spatial configuration was systematically varied every 3 min by shuffling the positions of the four corner stimuli across 24 configurations. **c**, Photograph of the 45 × 45 cm arena containing four salient corner landmarks, along with additional environment-specific landmarks, including a door and a holder. **d-e**, Example linear decoding of position along the *x*-axis **(d)** and y-axis **(e)** from population activity in mFC, dCA1, and lvOFC from the cups assay. Predicted position is shown in color and actual position in black. Scale bars, 30 cm (vertical), 1 minute (horizontal). **f-g**, Linear decoding error for position along the *x*- (left) and *y*-axes (right) in the cups assay **(f)** and social assay **(g)** across mFC, dCA1, and lvOFC. Each colored circle represents data from one mouse; horizontal lines and error bars indicate the mean ± SEM. Here and in subsequent figures, unless otherwise indicated, comparisons across three brain regions were assessed using a one-way ANOVA followed by pairwise Welch’s *t*-tests with Holm correction for multiple comparisons. For shuffle analyses, 100 shuffled controls were generated for each animal by circularly time-shifting calcium activity relative to position and evaluating each shifted recording across all cross-validation folds. Gray dots show the pooled shuffle-derived decoding errors, and gray horizontal lines indicate their means. For each brain region, the observed decoding error for each mouse was compared with the mean of that mouse’s shuffle distribution using a two-sided paired *t*-test across mice. (\**p <* 0.05, \*\**p <* 0.01, \*\*\**p <* 0.001, \*\*\*\**p <* 0.0001; n.s., not significant).

We have discovered a spatial code in the mouse medial frontal cortex (mFC) that exhibits this fundamental property of geometric maps. Linear regression of the animal’s *x* and *y* coordinates trained in one region of an arena can predict position in regions not used for training, a property we did not observe in hippocampal dorsal CA1 (dCA1) or lateroventral orbitofrontal cortex (lvOFC). This ability to extrapolate across physical space suggests that the mFC representation preserves spatial relationships in a form that may support abstract route planning and navigation.

### A spatial representation in mFC supports linear extrapolation across the environment

We used GCaMP6f and head-mounted miniscopes to record excitatory neurons in medial frontal cortex (mFC), hippocampal dorsal CA1 (dCA1), and lateroventral orbitofrontal cortex (lvOFC) as mice explored a square arena containing four salient corner landmarks, either four empty cages (cups assay) or four individually caged conspecifics (social assay) (Fig. 1b-c, Extended Data Fig. 1a-f). Activity was recorded across 24 sessions (3 min each), with the corner locations of conspecifics swapped between sessions (Figure 1b). This protocol supported sustained exploration (Extended Data Fig. 1g) and, while it was originally designed to study representations of social identity, it revealed a striking generalizable representation of space in mFC that was independent of social identity and became the focus of this study.

We explored neural representations of position along two orthogonal spatial axes. We performed linear regression independently for the *x* and *y* positions by reading out the activity recorded as animals explored the full arena. We tested the linear regression performance on held-out temporal blocks. Both *x* and *y* positions could be read out independently from mFC and dCA1 population activity, with markedly lower performance in lvOFC. In mFC and dCA1, the decoded position closely tracked the mouse’s actual position as it navigated the arena. This correspondence was weaker in lvOFC (Fig. 1d-e). To quantify decoding accuracy, we calculated the median absolute difference between the decoded and actual mouse positions along each axis (in cm) and compared it with a null distribution generated by circularly shifting neural activity relative to position. Decoding errors were significantly lower than those obtained from shuffled controls in mFC and dCA1, whereas the effect was weaker in lvOFC (Fig. 1f-g). Consistent with this, two-dimensional position could also be decoded using a nonlinear Bayesian decoder from both mFC and dCA1 populations, with greater decoding accuracy in dCA1. Decoding from lvOFC was substantially poorer (Extended Data Fig. 2).

Linear regression for *x* and *y* positions from the full arena does not establish that the underlying representation supports extrapolation across space. To test for extrapolation, we trained linear regression to predict either *x* or *y* position using activity from only one half of the arena. We then tested decoding accuracy on the other half, calculating the median absolute difference between the predicted and actual positions, which we refer to as generalization error. (Fig. 2b-d, Extended Data Fig. 3). Importantly, the linear regression model was constructed using only neural activity recorded when the animal was in the training half of the arena, without any information from the opposite half. For *x* position, linear regression was trained on the top half of the arena and tested on the bottom half, and vice versa; for *y* position, linear regression was trained on the left half and tested on the right half, and vice versa. Generalization error was averaged across the two train-test directions. As a control, we also trained and tested linear regression within the same half (Fig. 2a,c-d), which we refer to as decoding error.

**Fig. 2.**
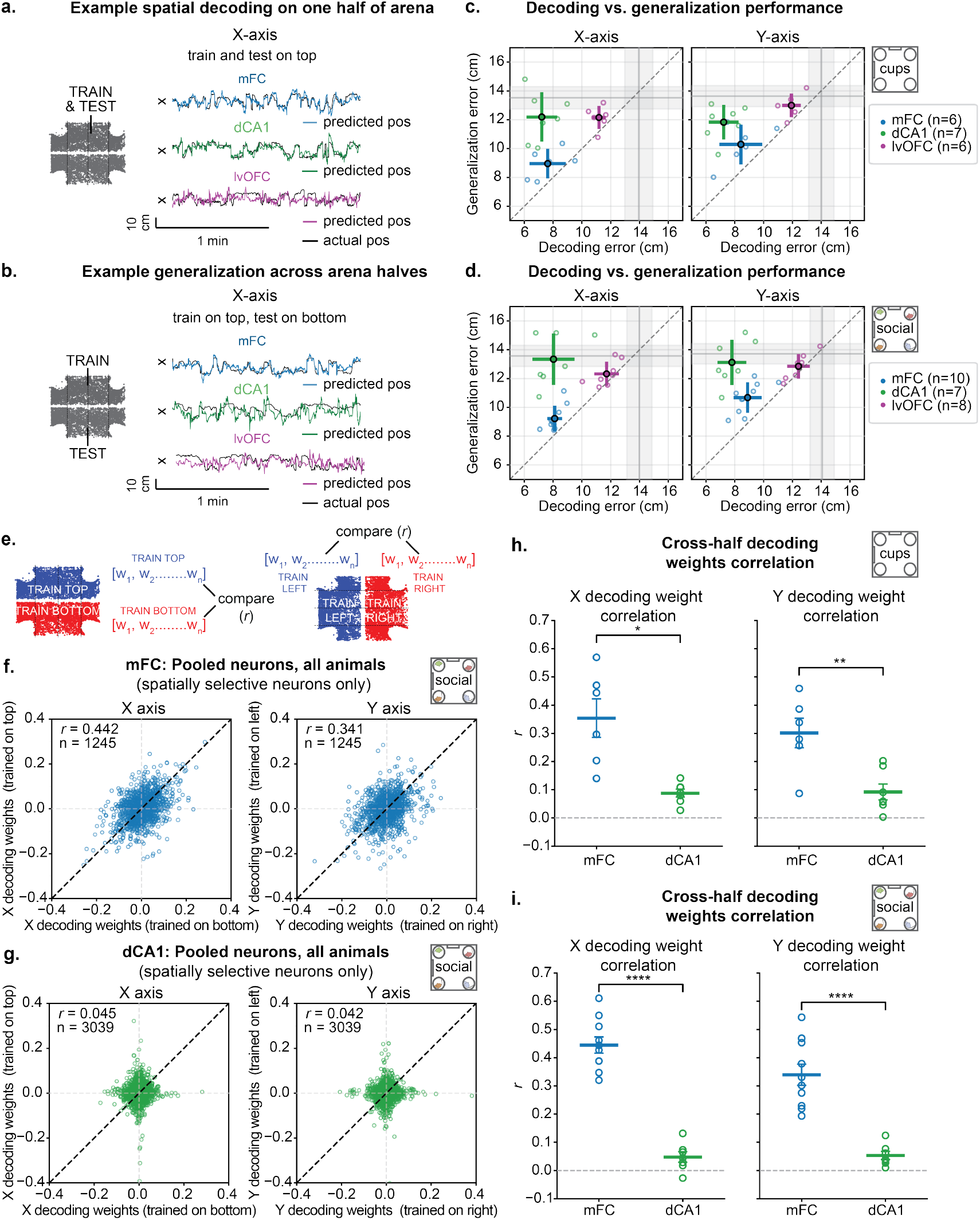
mFC supports linear extrapolation across space. **a-b**, Linear regression was trained to predict nose position along one spatial axis using population activity sampled from only one half of the arena along the orthogonal axis. For *x*-position, the arena was divided into top and bottom halves; for *y*-position, it was divided into left and right halves. Linear regression was tested **(a)** within the same arena half (decoding) or **(b)** on the other half (generalization), thereby testing prediction at positions along the orthogonal axis that were not represented in the training data. Examples show *x*-position in the social assay; both axes and both assays are quantified in **(c–d)**. Black traces show actual position; colored traces show predictions for each brain area. **c**, Decoding vs. generalization error across brain regions in the cups assay. For each mouse, generalization error was averaged across the two directions of cross-half testing, and decoding error was averaged across the two arena halves. Each circle represents data from one animal; filled points and error bars indicate the mean ± SEM **d**, Same as **(c)**, but for the social assay. **e**,. Schematic of the cross-half weight-correlation analysis. For *x*-position, independent linear regression models were trained on the top and bottom halves of the arena; for *y*-position, independent linear regression models were trained on the left and right halves. For each axis, the correlation between the weights assigned to individual neurons by the two half-specific linear regression models was calculated and compared between mFC and dCA1. **f-g**, Weights assigned to individual neurons by linear regression models trained on opposite halves are compared for *x*- (left) and *y*- (right) position for mFC **(f)** and dCA1 **(g)** populations in the social assay. Neurons were pooled across animals. For each animal and spatial axis, the two linear regression model weights for each neuron were divided by the Euclidean norm of all weights from both models. Each point represents one neuron. Pearson’s correlation coefficients (*r*) and numbers of neurons (*n*) are shown. We restricted the comparison of decoder weights across halves to spatially selective neurons, identified using a mutual-information test for position (see Methods). **h-i**, Correlation of cross-half decoding weights across animals in the cups assay (**h**) and social assay (**i**). Left, Pearson correlation coefficients between weights from the two *x*-position linear regression models. Right, Pearson correlation coefficients between weights from the two *y*-position linear regression models. Each circle represents one animal; horizontal lines and error bars indicate the mean ± SEM.

We observed that mFC representations extrapolate across space. The position predicted by linear regression trained on one half of the arena closely tracked the animal’s actual position in the opposite half, for both *x* (Fig. 2b) and *y* coordinates. In contrast, predictions from dCA1 populations deviated substantially from the animal’s actual position when tested on the opposite half (Fig. 2b). To quantify this difference, we compared generalization error with decoding error. In mFC, generalization errors remained low and were comparable to decoding errors. In contrast, dCA1 exhibited substantially higher generalization errors despite accurate within-half decoding, resulting in a much larger ratio of generalization to decoding error (Fig. 2c-d, Extended Data Fig. 3). Generalization remained robust when neuron numbers were matched across animals and brain regions (Extended Data Fig. 4) and across days in a subset of mFC recordings repeated on multiple days (Extended Data Fig. 5). In contrast, lvOFC performed poorly in both decoding and generalization (Fig. 2a-d, Extended Data Fig. 3).

Generalization performance in mFC was typically higher for *x* position than for *y* position, potentially reflecting asymmetries in environmental cues (Fig. 2c-d, Extended Data Fig. 3-4). Removing corner landmarks reduced generalization in mFC, whereas replacing conspecifics with empty cups had little effect (Extended Data Fig. 6), indicating that environmental landmarks are needed to shape and anchor the mFC spatial map.

The ability to extrapolate both the *x* and *y* coordinates that we observed in mFC implies that the neural displacement vectors across the two coordinate axes are approximately parallel in different parts of the arena. Any arbitrary movement in the 2D physical environment explored by the animal can be decomposed into displacements along these axes. As a consequence, arbitrary movements would be approximately represented by displacement vectors in neural activity space that do not depend on the animal’s initial position, a property that is important for planning and navigation, as discussed in the Introduction.

We next asked whether individual neurons contribute similarly to decoding a given spatial coordinate (i.e., have similar decoding weights) across opposite halves of the arena, and whether this consistency supports generalization in mFC. We trained linear regression models independently on different halves of the arena and compared the weights assigned to individual neurons in predicting *x* and *y* positions (Fig. 2e). We restricted the comparison of linear regression weights across halves to spatially selective neurons identified using a mutual-information criterion for position (8). Pooling neurons across all animals within each brain region, we found that linear regression weights were strongly correlated across opposite halves of the arena in mFC, whereas correlations were substantially weaker in dCA1 (Fig. 2f-g, Extended Data Fig. 7). In dCA1, we can clearly see that the distribution has the shape of a cross, indicating that the neurons important in the two halves belong to two non-overlapping populations. This difference was also evident when correlations were calculated separately for each animal, with significantly higher weight correlations in mFC than in dCA1 across the cups and social assays (Fig. 2h,i). These findings suggest that the contributions of individual mFC neurons to position decoding remain consistent across space, providing a basis for the generalization of the population code.

### Explaining extrapolation with neural models

What properties of mFC neuronal responses allow the population to extrapolate from explored to unexplored regions of space? The simplest way to construct a neural representation that generalizes across an environment is to have neuronal firing rates (*f*_*k*_ for neuron *k*) depend linearly on the *x* and *y* coordinates of the animal (*f*_*k*_ = *a*_*k*_*x* + *b*_*k*_*y*), where *a*_*k*_ and *b*_*k*_ are constants (Fig. 3a). The model’s ability to support extrapolation can be understood by examining the geometry of its neural encoding manifold. The neural activity space for *N* neurons is an *N*-dimensional space describing all possible population activity patterns. The encoding manifold is the subspace within this full activity space that describes how population activity varies as the animal occupies different locations in the environment. Ignoring noise, this manifold is a two-dimensional surface parameterized by the *x* and *y* coordinates of the animal’s position. For linear neuronal responses, this manifold is a flat two-dimensional surface with one direction encoding the *x* coordinate, and the other *y* (Fig. 3b). Because these two directions are independent, linear regression for *x* that is trained at a particular value of *y*, will inevitably generalize to any other value of *y*, allowing the kind of extrapolation we observed. The same argument applies to a *y* linear regression. However, as we show below, the mFC representation does not exhibit such a flat neural manifold. The key to the generalization ability of the linear model is that there is no nonlinear mixing of *x* and *y* dependencies in neuronal responses in the form of an *xy* term. To understand why such mixing disrupts generalization, consider linear regression trained at a particular *y* value, say *y*_1_, which will make use of both the pure *x* and the *xy*_1_ dependencies of the firing rates. At another *y* position, say *y*_2_, the second term will change to *xy*_2_, and thus linear regression will not generalize across *y* values. Here, for simplicity, we considered linear regression trained on a single value of *y*. However, the *xy* term remains disruptive even when training across a broader region, provided noise is present. Similar considerations apply to linear regression of *y*, which will also fail to extrapolate when mixed *xy* terms are present in the neural encoding. Indeed, adding a mixing term, *xy*, to the linear model described above destroys its ability to generalize (Extended Data Fig. 8). Nevertheless, there are certain nonlinear terms that can be added to the firing rates in the linear model while preserving generalization, as we will now show.

**Fig. 3.**
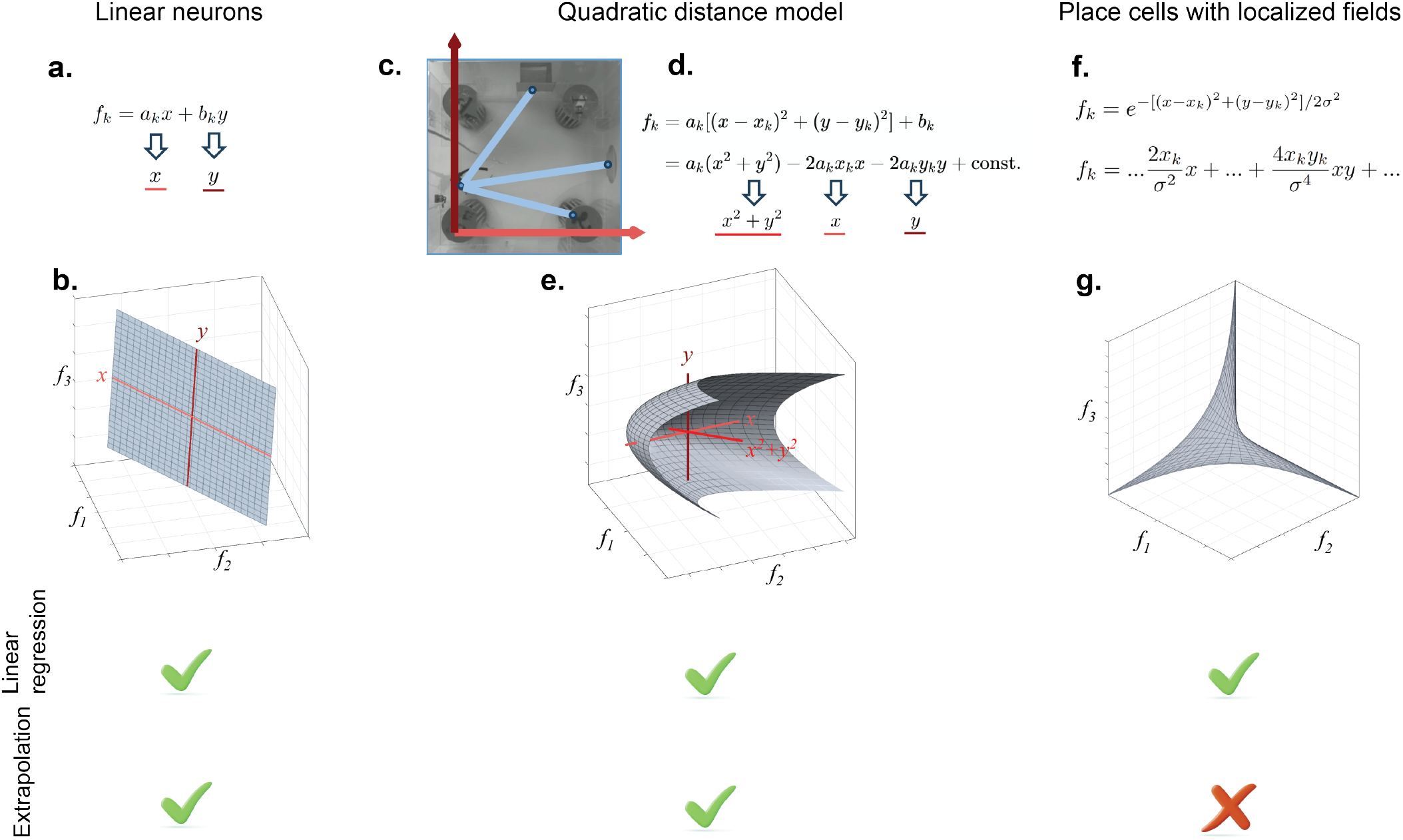
Idealized models of neural responses and their geometries. **a-b**, Linear model: the firing rate of each neuron is a linear function of the *x* and *y* coordinates of the animal. The manifold in neural activity space is a plane, making it possible to perform linear regression on both *x* and *y*. Linear regression can extrapolate. **c-e**, Quadratic distance model. Each neuron encodes the squared distance from a landmark. The neural manifold is a 3D paraboloid. In **(c)**, blue lines indicate distances from the animal’s position to different landmarks, while red arrows indicate the *x*- and *y*-axes. The red lines in **(e)** are the encoding directions for *x, y*, and *x*^2^ + *y*^2^. Linear regression can extrapolate. **f-g**, Place cells with localized fields. The neural responses are described as a Gaussian function. The field’s width is small relative to the environment. The manifold for three place cells is the highly curvilinear spiky representation in the plot. Linear regression does not generalize.

Recent work has shown that neurons in mFC encode distance to goals (9). Because distance calculation typically requires quadratic terms, we considered whether neurons with quadratic distance-dependent firing rates could give rise to a generalizable spatial map. Suppose that, when the animal is at location (*x, y*), neuron *k* in a neural population has a firing rate given by (Fig. 3c-d)

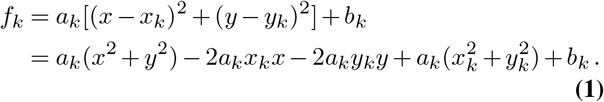

Here, *a*_*k*_ and *b*_*k*_ are parameters that determine the distance sensitivity and baseline rate of neuron *k*. The parameters *x*_*k*_ and *y*_*k*_ define the landmark point from which neuron *k* encodes the squared distance (*x* − *x*_*k*_)^2^ + (*y* − *y*_*k*_)^2^ (Fig. 3c). The second equality in the above equation expands the squares in the firing rate expression, revealing terms linear in *x* and *y* that enable linear regression for the two coordinates. This firing rate function, which depends on the squared distance from a landmark, avoids the mixing of *x* and *y*, such as the term *xy*, and hence allows for extrapolation.

In the squared distance scheme, the landmark points (*x*_*k*_, *y*_*k*_) could be different for each neuron, or multiple neurons could share the same landmark. The only requirement is that there must be at least three different, non-collinear landmarks (see Methods). We can construct the neural coding manifold for a population of neurons with squared-distance tuning by examining the various terms in the rightmost expression of equation 1. The last of these terms, 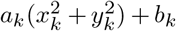, is just a constant. The other three terms define three directions in the neural activity space along which the quantities *x*^2^ + *y*^2^, *x*, and *y* are represented. The coordinates of these vectors are *a*_*k*_ for *x*^2^ + *y*^2^, *a*_*k*_*x*_*k*_ for *x* and *a*_*k*_*y*_*k*_ for *y*. Although they span three dimensions, these vectors are not necessarily orthogonal. Because of the combined linear and quadratic dependence of the firing rates on the spatial coordinates (*x, y*), the coding manifold for this squared distance model is a three-dimensional paraboloid in neural activity space (Fig 3e).

In the absence of noise, the *x*^2^ + *y*^2^, *x*, and *y* directions define a three-dimensional subspace. When performing principal component analysis, we would find three dominant PCs that span this space. These PCs are linear combinations of the three vectors defining the three directions. This ensures that the top PCs have a linear dependence on *x* and *y* built into them, which supports linear decoding of these coordinates. The model thus makes two predictions (see also Methods): there should be 3 dominant PCs that correspond to linear combinations of the *x*^2^ + *y*^2^, *x*, and *y* directions, and linear, generalizable decoding should be based solely on them.

We test these predictions against the data in the following section.

How does a standard place-cell representation compare to the squared distance coding we have introduced? The firing rate of a place cell (labeled by an index *k*) with a field of width *σ*_*k*_ centered at the point (*x*_*k*_, *y*_*k*_) can be modeled as a Gaussian function of the animal’s position (Fig. 3f),

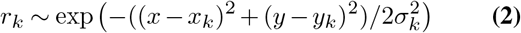

Although this firing rate depends on the squared distance from a landmark (the place-field center), just as in the squared distance model, the presence of the exponential function in this case introduces a large number of mixing terms, such as *xy*, that destroy generalization. The neural manifold of a population of such place cells has the spiky shape illustrated in Fig. 3g. Three place cells with firing rates defining the coordinates in this three-dimensional figure peak at different locations. If the animal moves from one of these locations to the other, one neuron will be silenced and another activated. This makes the coding manifold in this reduced space three-dimensional. Including a larger population of neurons generates a manifold with a dimensionality that grows in proportion to the number of neurons in the neural population with non-overlapping place fields. By contrast, the squared distance representation is low-dimensional regardless of the number of neurons. Nevertheless, for the place cell representation, there are independent directions that linearly encode *x* and *y*. Thus, place cell encoding can support linearly decodable coordinates, but such linear regression decoders cannot generalize because of the presence of *xy* and higher-order mixing terms in the firing rates. Rather than the low-dimensional decoding predicted by the squared distance model, the place cell model predicts that a larger number of PCs encode *x* and *y*.

We note that for very large place fields, the exponential tuning function can be approximated by a quadratic function of position, which may allow them to generalize. Indeed, if *σ*^2^ ≫ (*x* − *x*_*k*_)^2^ + (*y* − *y*_*k*_)^2^ for all the points in an environment, expression 2 reduces to *f*_*k*_ ~ (1 − ((*x* − *x*_*k*_)^2^ + (*y* − *y*_*k*_)^2^)*/*2*σ*^2^), which is of the same form as the squared distance encoding in equation 1. However, the large place-field model of generalization faces a tradeoff. The quadratic approximation is only valid when ((*x*−*x*_*k*_)^2^ +(*y* − *y*_*k*_)^2^)*/*2*σ*^2^ ≪ 1. If place fields are not sufficiently large, higher-order terms become important and the quadratic approximation breaks down. Conversely, if place fields are much larger than the arena, their spatial modulation becomes small, making the representation sensitive to noise. Thus, for large place fields to support generalization, they must be large enough for the quadratic approximation to hold, but not so large that spatial modulation becomes negligible.

### A small number of dimensions encode position in mFC but not in dCA1

The squared distance model predicts a low-dimensional manifold in which the *x* and *y* coordinates are encoded in only 3 dimensions, each corresponding to a linear combination of *x, y*, and *x*^2^ + *y*^2^. In contrast, the place cell model predicts that the *x* and *y* coordinates are represented in a high-dimensional subspace. We now show that mFC is better described by the squared distance model and dCA1 by the place cell model.

We performed principal component analysis (PCA) on spatial rate maps computed across the full arena. For each neuron, we computed an 8 × 8 spatial rate map and vectorized activity across spatial bins. Because the four corner bins containing the landmarks were not sampled, this yielded 60 spatial bins. The variance explained by individual PCs declined more rapidly in mFC than in dCA1, and the participation ratio of the explained variance was significantly lower in mFC across both assays (Extended Data Fig. 9), indicating that spatial population activity in mFC was lower-dimensional than in dCA1.

We then used the weight vectors obtained from the earlier linear regression analysis (Fig. 2) to identify the *x* and *y* encoding directions. For the *x*-position, these linear regression models were trained separately on the upper and lower halves of the arena, and for the *y*-position, on the left and right halves. We computed the absolute cosine similarity between each PC and each linear regression weight vector separately. In mFC, the encoding directions for *x* and *y* aligned primarily with the top three PCs (Fig. 4a,e). Specifically, the *x* encoding direction showed the greatest cosine similarity with PCs 1 and 2, whereas the *y* encoding direction showed strong alignment with PC 3. In dCA1, on the other hand, the encoding directions were more evenly aligned with multiple PCs (Fig. 4b,f). To quantify this difference, we computed the participation ratio of the cosine similarities between each encoding direction and the PCs (Fig. 4c,g). We found that the number of dimensions contributing to position decoding was significantly lower in mFC than in dCA1.

**Fig. 4.**
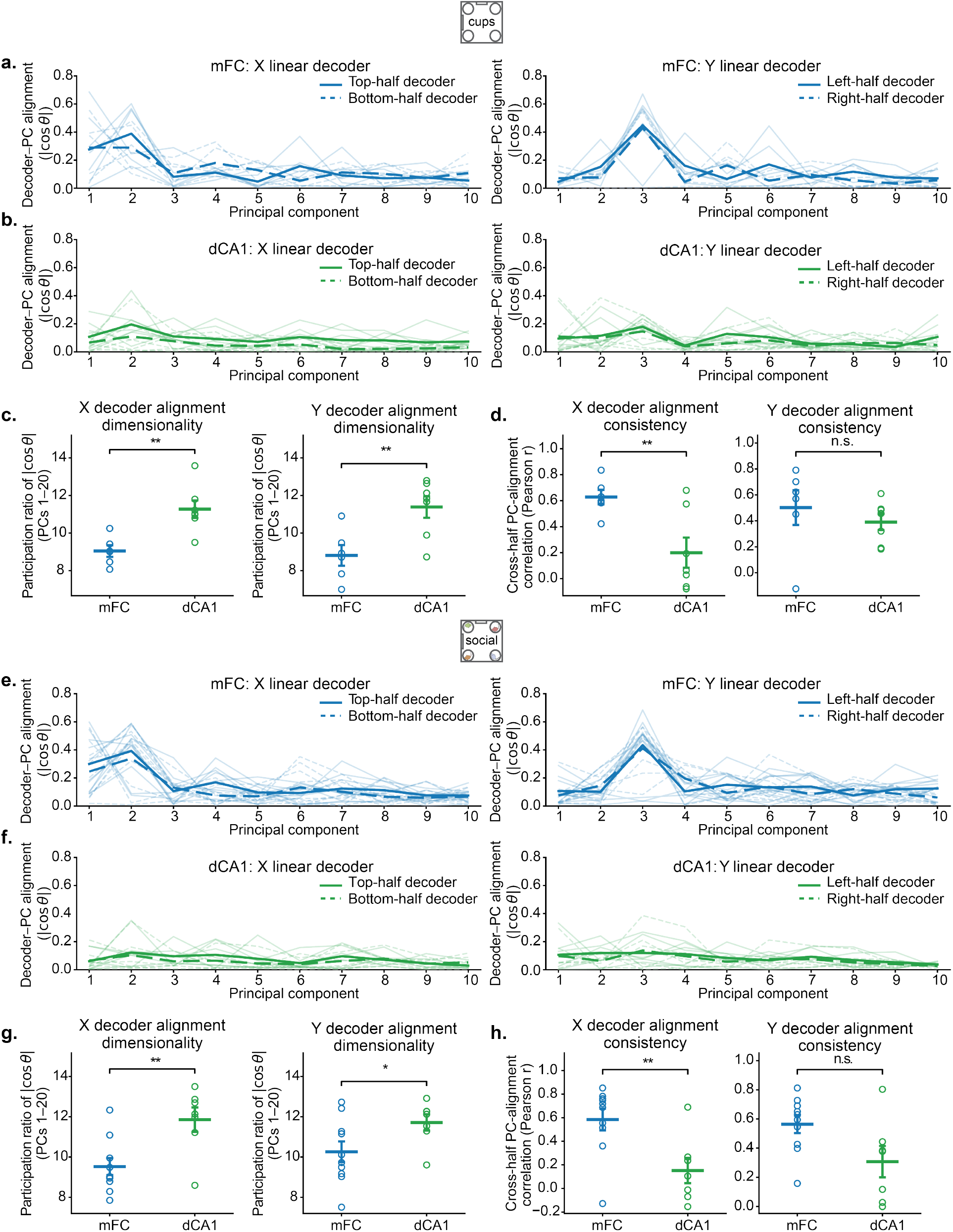
Spatial decoding axes are concentrated in fewer principal components and are more consistent across space in mFC than dCA1. **a-b**, Alignment between whole-arena principal components (PCs) and linear regression axes for *x*-position (left) and *y*-position (right) in the cups assay for mFC **(a)** and dCA1 **(b)**. For *x*-position, linear regression models were trained separately on the top and bottom halves of the arena; for *y*-position, linear regression models were trained separately on the left and right halves. For each linear regression model, the alignment with every PC was quantified as the absolute cosine similarity between the PC loading vector and the linear regression weight vector. Faint lines indicate individual mice, and thick lines indicate the mean across mice. **c**, Participation ratio of the cosine similarities between each linear regression weight vector and the PCs, quantifying how broadly each decoding axis was distributed across principal components. Lower values indicate that a lower number of PCs that are aligned to the *x* and *y* encoding directions. **d**, Consistency of encoding directions-PC alignment profiles across opposite halves of the arena, quantified as the Pearson correlation (*r*) between the two alignment profiles for *x*- and *y* coding directions. **e-h**, Same analyses as in a-d for the social assay. Circles in c,d,g,h indicate individual mice; horizontal bars show mean ± S.E.M.

Lastly, we asked whether the alignment profile between the coordinate encoding directions and PCs was stable across the two halves of the arena used to train the linear regression models. We computed the Pearson’s correlation coefficient between the alignment profiles of linear regression models trained on opposite halves of the arena and found significantly higher correlations in mFC than in dCA1 (Fig. 4d,h). Together, these results suggest that linear regression in mFC relies on a low-dimensional neural subspace that remains stable across different parts of the environment, providing a basis for generalization.

Next, we tested the squared distance model prediction that the three dominant spatial patterns of population activity should correspond to linear combinations of *x*^2^ + *y*^2^, *x*, and *y*. We performed singular value decomposition (SVD) on the population spatial rate maps and visualized the first three spatial modes by reshaping the corresponding spatial scores into two-dimensional maps. Like PCA, SVD identifies dominant patterns of population activity. In this analysis, each spatial mode can be interpreted as a weighted sum of the population-mean-centered spatial activity maps of individual neurons. Next, we fit each spatial mode with the most general quadratic model *f* (*x, y*) = *a*(*x*^2^ + *y*^2^) + *bxy* + *cx* + *dy* + *e* to quantify the relative contribution of each term to its spatial structure. This analysis revealed a striking spatial organization in mFC (Fig. 5a-c, Extended Data Fig. 10). Mode1 and Mode2 could be described as a linear superposition of *x*^2^ + *y*^2^ and *x*, with a smaller contribution from *y*, and Mode3 was dominated by *y*, with some contribution from *x*. The contribution of the *xy* term to all three modes was negligible. Thus, the spatial structure of the first three modes in mFC was consistent with the first prediction of the squared distance model. In contrast, the coefficients for the *xy* term in the models fit to dCA1’s spatial modes were, on average, more than 6 times larger than for mFC (Extended Data Fig. 11). This is one of the differences that explains why mFC can generalize but dCA1 cannot.

**Fig. 5.**
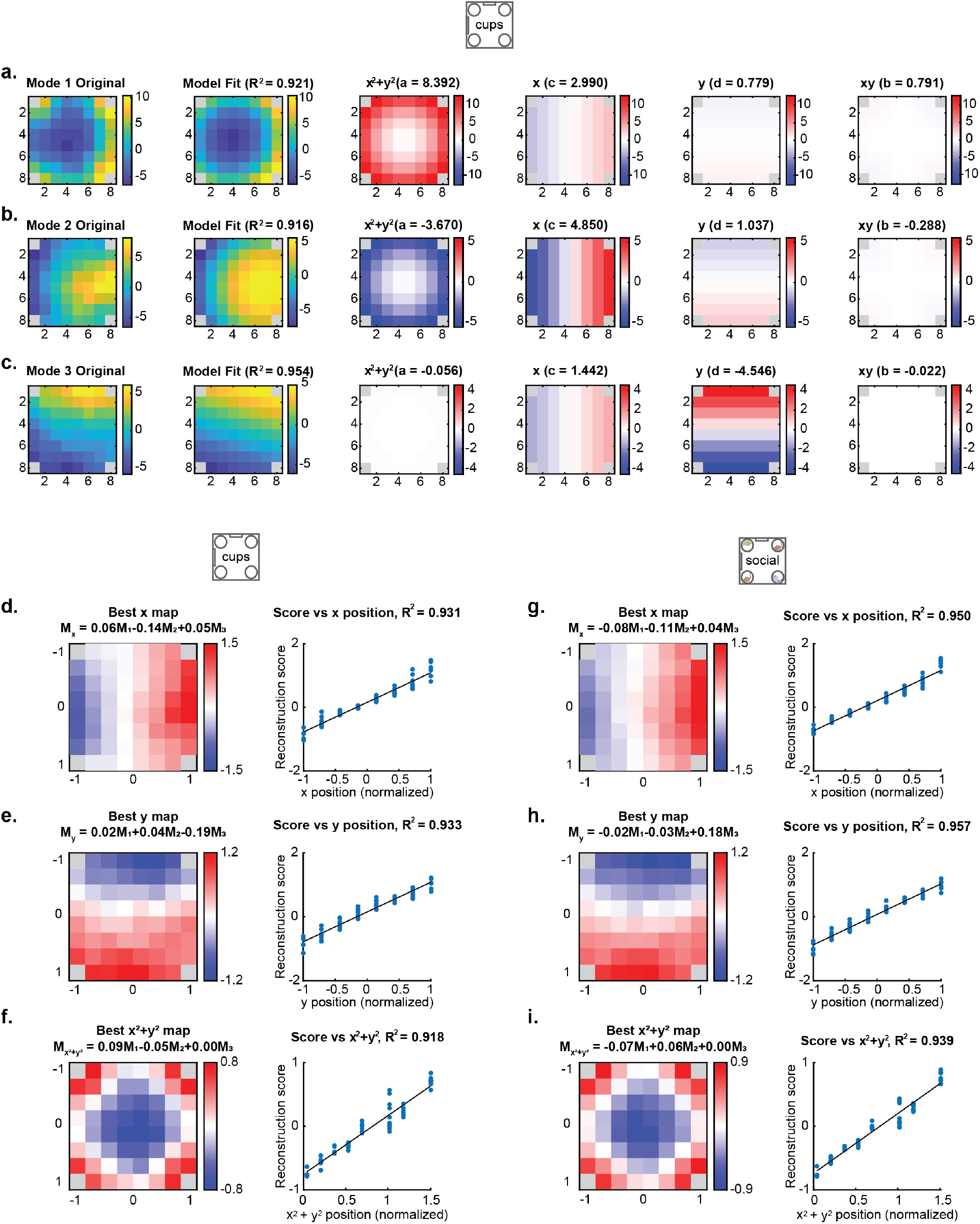
The leading spatial patterns of neural activity identified by SVD are explained by a squared distance-based model in mFC. **a-c**, The first three spatial modes obtained by singular value decomposition (SVD) of mFC population activity in the cups assay. For each mode, the original spatial score map (left) is fit with the model *f* (*x, y*) = *a*(*x*^2^ + *y*^2^) + *bxy* + *cx* + *dy* + *e*, using spatial coordinates normalized to [−1, 1]. The fitted map and *R*^2^ are shown alongside the individual contributions of the *x*^2^ + *y*^2^, *x, y*, and *xy* terms; the corresponding fitted coefficients are indicated above each component map. **d-f**, Linear combinations of the first three spatial modes were fit by ordinary least squares to reconstruct *x* position **(d)**, *y* position **(e)**, and *x*^2^ + *y*^2^ **(f)** in the cups assay. Left, reconstructed spatial map; right, reconstruction score as a function of the corresponding spatial variable. Black lines show linear fits; *R*^2^ values indicate reconstruction performance. **g-i**, Same analysis as in d-f for the social assay, reconstructing x position **(g)**, y position **(h)**, and *x*^2^ + *y*^2^ **(i)**. Gray corner bins were excluded from all analyses.

We further tested whether the low-dimensional spatial representation in mFC supports linear readout of position across different parts of the environment by asking whether the first three spatial modes were sufficient to reconstruct *x* and *y* positions. We computed the linear combinations of the first three spatial modes that optimally reconstructed *x, y*, and *x*^2^ + *y*^2^ (Fig. 5d-i). All three reconstructions achieved high *R*^2^ values, and the reconstruction score as a function of the respective spatial term was consistent across the entire arena. In sum, our PCA and SVD analyses revealed that mFC satisfies the two predictions of the quadratic distance model, whereas dCA1 satisfies those of the place cell model.

#### The activity of single neurons supports a quadratic distance model

Equation 1 describing individual neural responses in the squared distance model is clearly an idealization. Neural responses are typically very diverse and often seemingly disorganized (10, 11). Neurons respond nonlinearly to multiple task-relevant variables (mixed selectivity), especially in cognitive brain areas such as the prefrontal cortex (12). The hippocampus and entorhinal cortex are no exception: the representations in both CA1 and dentate gyrus are highly distributed, and the individual responses are very diverse (8). Analogously, even selected grid cells exhibit mixed selectivity and are highly heterogeneous (13). Because of this diversity of responses, it is difficult to draw conclusions from analyzing firing patterns of individual neurons, and this is also the case in our recordings of mFC.

The key results from our idealized model are that *x*^2^ + *y*^2^, *x*, and *y* should be strongly represented across the neural population, which will happen if the firing rates of a sufficient number of cells are at least partially described by equation 1. In other words, there can be additional terms in the expressions for the firing patterns of individual neurons, but the squared distance model should account for a large fraction of the total responses across the population. To investigate this, we examined how well equation 1 describes the firing patterns of the neurons we recorded.

We fit the squared distance model to the neuronal spatial fields in mFC and dCA1. Supporting the model’s hypothesis, we found that a substantially larger fraction of mFC neurons was well described by the quadratic distance model than in dCA1 across multiple RMSE thresholds and in both the cups and social assays (Fig. 6a). We next asked whether neurons that were well described by the squared distance model contributed more consistently to position decoding across different parts of the arena. As in our earlier analysis, we used linear regression trained separately on opposite halves of the arena and compared the weights assigned to individual neurons by each linear regression model. We found that mFC neurons that were well described by the model had more consistent linear regression weights across the two halves than neurons that were less well fit. This was true for *x*-position linear regression trained on the upper and lower halves (Fig. 6b, Extended Data Fig. 12a) and for *y*-position linear regression trained on the left and right halves (Fig. 6c, Extended Data Fig. 12b). These results suggest that neurons well described by the squared distance model contribute more consistently to position decoding across space, supporting their role in generalization.

**Fig. 6.**
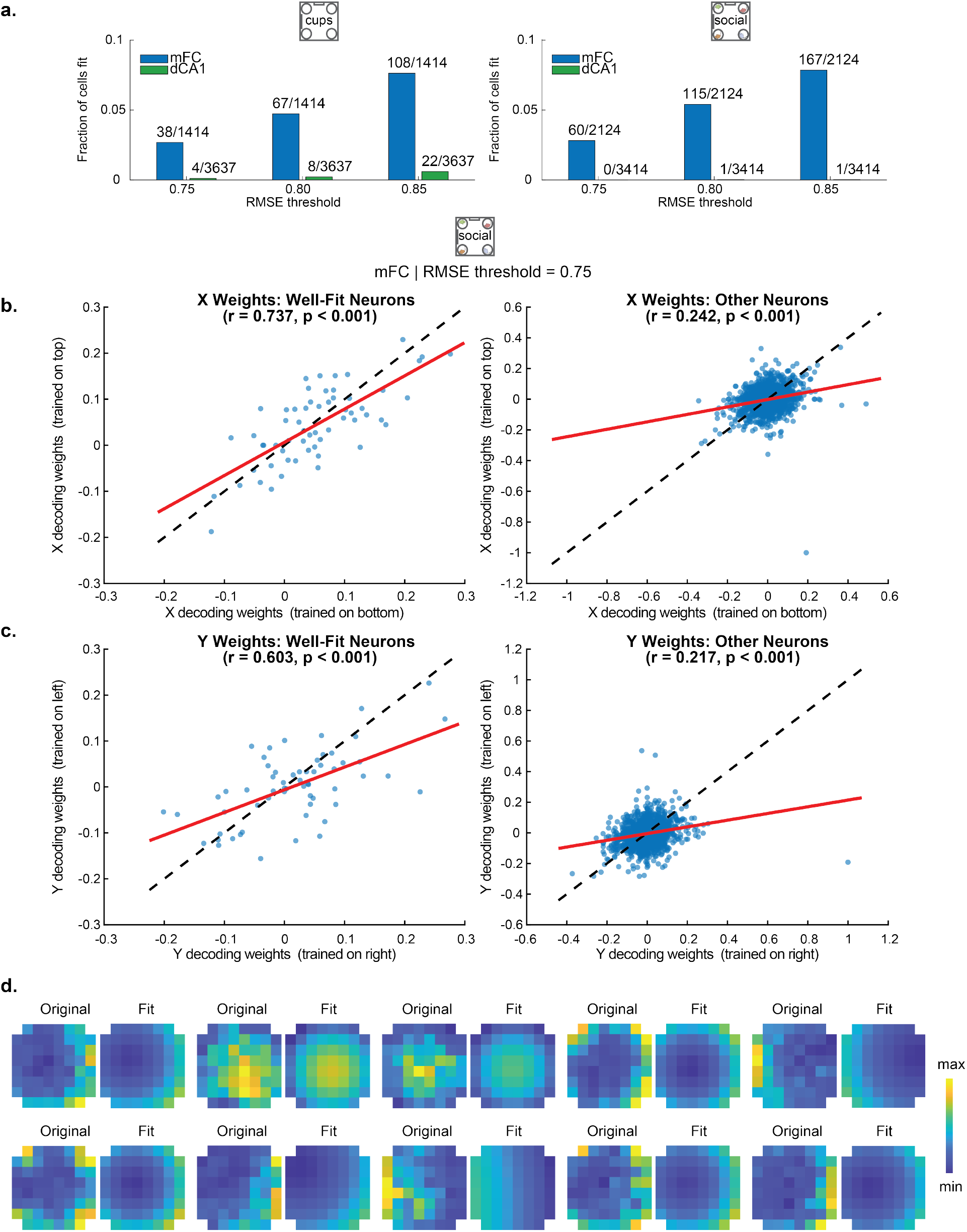
A subset of mFC neurons is well described by a squared distance-based spatial model. **a**, Fraction of neurons in mFC (blue) and dCA1 (green) classified as well fit by the squared distance model at RMSE thresholds of 0.75, 0.80 and 0.85 of the standard deviation of the spatial bins in the raw rate map, for the cups (left) and social (right) assays. **b**, Normalized *x*-position linear regression weights obtained from the top versus bottom halves of the arena in the social assay, shown separately for neurons well fit by the squared distance model (left) and all other neurons (right), using an RMSE threshold of 0.75. Each point represents one neuron. Red lines indicate linear fits and dashed lines indicate the identity line; Pearson’s *r* and corresponding *p* values are shown above each plot. **c**, Same as b, but for *y*-position linear regression weights obtained from the left versus right halves of the arena. **d**, Ten representative mFC neurons classified as well fit from the cups and social assays at an RMSE threshold of 0.75, showing the observed spatial activity map (Original) and its corresponding distance model fit (Fit). For each neuron, the original and fitted maps are displayed using a common color scale spanning the minimum to maximum value across the two maps; the color bar indicates this normalized minimum-to-maximum range.

The spatial fields of neurons well described by the squared distance model were nevertheless diverse, with a broad distribution of parameters *a*_*k*_, *b*_*k*_, *x*_*k*_, and *y*_*k*_ (Fig. 6d). There are cells with *a*_*k*_ *<* 0 that look somewhat like large place cells. Some cells with *a*_*k*_ *>* 0 exhibit strong activation near the edges of the arena, similar to border cells (14). Other cells could also be interpreted as boundary vector cells (15), as seen in the subiculum and entorhinal cortex, or object vector cells (16), as seen in MEC, which fire when an animal is at a specific distance and allocentric direction relative to a boundary or object. Nevertheless, this diverse population of neurons, each at least partially encoding squared distance from different landmarks, can collectively form a low-dimensional spatial manifold capable of extrapolation.

## Discussion

We identified a new spatial code in mFC that possesses a property critical for planning and navigation. This representation encodes physical displacement in a manner invariant to the starting point, enabling predictions of future locations along arbitrary routes in unexplored environments. An arbitrary route can be decomposed into a sequence of displacements, each represented by a vector connecting two successive positions. When using a map in an atlas, we can predict our final location after a series of movements (e.g., move East, North, East, South) by simply adding these displacement vectors to our initial position vector. This additivity property also holds for the *x,y* subspace of mFC neural maps—a coordinate system that can be inferred by exploring a restricted portion of the environment. This principle does not apply to idealized place cells with localized fields. For place cells, the neural representation of a displacement depends on its location within the environment. Two identical physical movements (such as equal eastward displacements from different starting locations) evoke distinct changes in population activity, each silencing the cell active at the starting location and activating a different cell encoding the final location.

In mFC, we showed that parallel physical movements along one of the two coordinate axes are represented by parallel vectors in neural activity space. This property manifests itself in the ability to cross-generalize (17): a decoder trained to extract the animal’s *x* and *y* coordinates in one region of the arena successfully generalizes to other regions. This is possible only because the *x* and *y* encoding directions in neural activity space are independent and remain approximately constant throughout the environment. While the *x* and *y* axes represent just two orthogonal directions in physical space, any arbitrary displacement vector can be formed by a linear combination of vectors aligned to the axes. Consequently, the highly desirable property of starting-point invariance extends to any arbitrary displacement in the environment. Ultimately, this reveals that the mFC spatial code is fundamentally compositional: every route can be decomposed into a series of displacements, and the location reached after each displacement can be predicted by simply adding the corresponding vectors in neural activity space. Compositionality is fundamentally important for achieving difficult forms of generalization—a computational principle essential not only for flexible navigation but also for abstract reasoning and cognition more broadly.

The neural maps in the hippocampus, MEC and mFC, are markedly different, with distinct computational properties, which may result in different functions in spatial navigation. In all three cases, the coding manifold is curvilinear, but it is high-dimensional in dCA1 and relatively low-dimensional in both mFC (paraboloid) and EC (torus, at least when the analysis is restricted to grid cells (18)). What are the computational implications of these different geometries?

It is possible to extract the animals’ *x* and *y* coordinates by reading out place cells. To see why, consider a population of neurons with highly localized place fields. By computing the weighted sum of the neurons, we can readily obtain the animal’s *x* coordinate, provided each cell’s weight scales with the coordinate of its peak firing rate. In dCA1, in contrast to mFC, linear regression must be trained over the entire arena. Because the neural code varies across the arena, with different place cells activated in different regions, linear regression does not generalize, as shown by our analysis of dCA1 activity. This high-dimensional neural code of space, however, has an advantage, not necessarily for navigation, but for memory. In artificial neural networks, the memory capacity scales with the dimensionality of the neural representations (19).

Grid cells share some properties with the mFC representation we have identified. Because the grid cell pattern repeats, activity in an unexplored region can be predicted from observed grid cell activity once the phase, orientation, and periodicity of a grid cell have been determined. Furthermore, if, as is widely believed, MEC integrates velocity signals that shift the grid pattern (20), a displacement in an unknown area will produce the same phase change as in a known area. However, translating a representation by integration is not ideal for planning long trips, because it requires simulating them mentally.

Grid cells provide a highly efficient, combinatorial spatial code. By leveraging multiple modules with distinct spatial periods, the grid system can represent very large environments without sacrificing spatial resolution (21). However, extracting spatial coordinates *x* and *y* from the grid code is challenging using simple linear regression. Position information can be accurately recovered only if the multi-scale grid modules are processed through a non-linear decoder. For instance, the grid code can be non-linearly transformed into localized place cell responses (22), from which position can subsequently be read out linearly.

In mFC, due to quadratic distance scaling, firing rates can easily saturate, limiting the coding range in large environments. One possible solution is the existence of multiple mFC maps operating at different inherent spatial scales. Alternatively, the spatial scales may be dynamically adjusted to fit the environment. A direct analog of this dynamic rescaling is found in experiments on time encoding, where the time elapsed since a relevant event is encoded by ramping neural activity. This activity peaks at the next relevant event. Interestingly, when the time interval between two events increases, the slope of the ramp is dynamically adjusted so that it always reaches the same peak at the next relevant event (23–25). Similar mechanisms could stretch or compress the mFC spatial map depending on the current environment’s physical boundaries.

The hippocampus plays an important role in learning and memory, whereas it is less clear whether it is important for navigation in familiar environments. Rodent experiments using the Morris water maze established that while hippocampal lesions severely impair the learning of new hidden platform locations, animals trained extensively prior to surgery retain intact spatial navigation capabilities, relying on consolidated remote spatial representations (26, 27). Likewise, studies of amnesic patients demonstrate that remote spatial representations acquired decades before hippocampal damage remain intact, enabling accurate mental navigation and vector planning through childhood neighborhoods (28).

MEC lesions severely impaired the acquisition of new spatial platform locations in the Morris water maze (29). Animals with MEC lesions exhibited learning deficits comparable to those with complete hippocampal damage, but could achieve good spatial performance in highly familiar environments. However, sometimes their underlying behavioral flexibility is impaired. When tested on novel starting configurations or flexible memory demands, animals without MEC input show persistent navigation deficits (29). In familiar openfield navigation, MEC-lesioned animals show impairments in updating their trajectories when distal landmarks are altered, indicating an inability to flexibly adapt path integration to visual landmarks (30). Causal manipulation studies demonstrate that while mFC inactivation leaves simple locomotion and familiar spatial memory intact, it severely impairs goal-directed planning, flexible route switching, and prospective trajectory generation in complex or dynamically changing environments (31–35).

We showed that linear regression on the animal’s coordinates successfully generalizes to an area of the arena excluded from the decoder’s training set. This is a difficult form of out-of-distribution generalization that necessitates highly structured neural representations. For linear generalization to succeed, the coding directions for *x* must remain parallel across all values of *y*. In a high-dimensional state space, such parallelism is exceptionally rare, as random vectors are nearly orthogonal. When representations encode *x* and *y* along independent, parallel axes, they are referred to as “disentangled” in the machine learning literature and are important for robust generalization. In neuroscience, we have previously termed these representations “abstract,” because they contain low-dimensional subspaces dedicated exclusively to single variables (e.g., an axis that encodes *x* while remaining invariant to *y*). Such abstract representations have been observed across diverse species and brain regions, including the dorsolateral prefrontal cortex and hippocampus in macaques (17), the hippocampus in humans (36), inferotemporal cortex (37), sensory cortices (38), area CA2 (39) and mFC in navigation tasks(40) in rodents. Crucially, the presence of these geometric structures is correlated with the capacity for cognitive inference (36). How these disentangled representations naturally form remains a significant, unresolved challenge in machine learning (41). In our study, the latent variables are the relatively simple spatial coordinates, *x* and *y*. If our quadratic distance model holds, these disentangled subspaces do not require complex learning rules; they emerge naturally simply by encoding the squared Euclidean distances to multiple environmental landmarks (Methods).

The neural maps that we observed in the frontal cortex may provide a basis for vector computation in navigation and complex route planning. In addition, given the simplicity of the quadratic distance model, the same principles might apply more broadly to any cognitive map in which “navigation” in a non-spatial latent space describes a general form of abstract reasoning.

## Methods

### Mice

All procedures were performed in accordance with the U.S. NIH Guide for the Care and Use of Laboratory Animals and were approved by the Institutional Animal Care and Use Committee at Columbia University. Adult male C57BL/6J mice (Jackson Laboratory) were used as subject mice, beginning at 12-25 weeks of age. Animals were housed either individually or, whenever possible, in groups with littermates (2-5 per cage). Stimulus mice were age- and sex-matched and were primarily of the C57BL/6J strain. In a subset of experiments imaging medial frontal cortex (mFC) (2 of 10 subject mice), two stimulus mice were C57BL/6J and two were CD1; as no differences were observed, data were pooled for analysis. Mice had ad libitum access to food and water and were maintained on a 12-hour light/dark cycle.

### Stereotactic surgery

To enable calcium imaging, stereotactic viral injections were performed in each target region in separate cohorts of mice followed by implantation of a gradient refractive index (GRIN) lens above the injection site. Injection coordinates (relative to bregma) were: mFC (ML: ±0.3 mm, AP: +2.0 mm, DV: −1.8 mm), dCA1 (ML: +1.4 mm, AP: −1.9 mm, DV: −1.35 mm), and lvOFC (ML: −1.0 mm, AP: +2.4 mm, DV: −2.45 mm). Mice were anesthetized with isoflurane and administered analgesia (buprenorphine XR, 3.25 mg/kg, subcutaneous) for all surgical procedures. The scalp was shaved and mice were positioned in a stereotactic frame. Eyes were protected with ophthalmic ointment (Patterson Veterinary) to prevent corneal drying. Body temperature was maintained using a heating pad with a feedback controller. Lidocaine was applied topically to the scalp to locally anesthetize the incision site. The skull was exposed and cleaned using sterile cotton swabs. A machine screw was secured to the skull to serve as an anchor for the dental cement. Prior to injection, a circular craniotomy (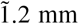 diameter) was made over mFC and lvOFC. For these regions, overlying tissue was gently aspirated to minimize tissue pressure prior to viral injection. For dCA1, a burr hole was made at the target site prior to injection. Injections were performed using a Nanoject III (Drummond Scientific) with a pulled glass micropipette. A total volume of 500 nL of AAV1.CaMKIIa.GCaMP6f.WPRE.bGHpA (Inscopix) was delivered in 10 nL pulses (10 or 23 nL/s) with a 10 s inter-pulse interval for 50 cycles into mFC or lvOFC (3.35 × 10^12^ vg/mL) or dCA1 (1.34 × 10^12^vg/mL). Injections into mFC were performed in either hemisphere, given its proximity to the midline, whereas dCA1 and lvOFC injections were performed in the right and left hemispheres, respectively. The pipette was left in place for 8-10 min following injection for mFC and lvOFC, and for 25 min for dCA1, before slow retraction. For dCA1, the craniotomy and aspiration were performed after viral delivery. A GRIN lens (1.0 mm diameter, 4.0 mm length; Inscopix) with an integrated baseplate was implanted above the craniotomy and centered over the injection site. The lens was secured using Metabond (Parkell), and the baseplate assembly was further stabilized and covered with black dental cement (Ortho-Jet). After 6-8 weeks, the level of GCaMP6f expression was evaluated using the miniscope. Imaging experiments began once viral expression was deemed sufficient based on visual inspection.

### Histology

Mice were deeply anesthetized with ketamine/xylazine and transcardially perfused with 4% paraformaldehyde (PFA). Heads with implants were collected and post-fixed in PFA for 24-48 h, after which brains were extracted and further post-fixed for 24 h. Coronal sections (100 µm) were prepared using a vibratome (Leica). Sections were stained with far-red NeuroTrace (640/660; Thermo Fisher Scientific) to label neuronal cell bodies. Images were acquired using a fluorescence microscope (Nikon AZ100 Multizoom) equipped with a motorized stage and automated acquisition software for high-throughput imaging. Histological analyses were performed to verify GRIN lens placement and to assess GCaMP expression based on native fluorescence.

### Behavioral assay

#### Arena design

Male subject mice explored a 45 × 45 cm square arena with an open top (Noldus). The arena was positioned on a table within a three-walled enclosure, also open at the top. A commutator and overhead camera were mounted above the arena, with cables routed from the ceiling into the enclosure. One wall of the arena contained a door through which the experimenter introduced the subject mouse and repositioned stimuli (empty cages or caged conspecifics, see below) at 3-min intervals during the experiment. For consistency, spatial references (left/right, upper/lower) were defined relative to this door: the wall containing the door was designated as the “left” side of the arena. The wall perpendicular to the door wall (to the left when facing the door) was defined as the “upper” wall and contained an empty holder as a landmark (see Fig. 1c). The arena walls were transparent, allowing animals visual access to distal cues in the surrounding room.

#### Assay structure

We performed three behavioral assays during the light phase of the 12 h light-dark cycle. In the first (empty arena), animals freely explored the arena for 30 min. Exploration was not sustained beyond this duration, with animals exhibiting limited movement. The remaining assays used a four-chamber paradigm. In the cups assay, four empty carousel cages (10 cm diameter, 20 cm height; acrylic with gray top; Noldus) were placed at the four corners of the arena. The positions of the cages were shuffled every 3 min such that all 24 possible permutations were presented to the animal in a pseudo-random sequence. In the social assay, four novel male mice were placed within the corner cages, and the same shuffling protocol was applied. For mFC recordings, animals experienced these assays in varying orders across days, and some experienced subsets of assays across multiple days. In addition, a subset of animals was tested in related assays (e.g., with one or two conspecifics in the same chamber), as the experiments were originally designed to probe social-spatial behavior. Decoding and generalization results were consistent across assay orders, both when pooling across all animals and experimental days and when analyzing cohorts with matched assay sequences separately. For analyses presented in Fig. 2, we used the first experimental day on which each mFC-implanted animal experienced the given assay (cups or social) in cases where animals were tested on multiple days. For recordings in lvOFC and dCA1, most animals followed a fixed sequence: empty arena (day 1), cups assay (day 2), and social assay (day 3). Two lvOFC- and three mFC-implanted animals experienced only the social assay, and two dCA1 animals were tested in a variable assay order similar to that used for mFC mice. Overall results were comparable across conditions. Entire recording sessions (i.e., full assay) with fewer than 50 identifiable neurons or insufficient exploration (defined as having fewer than 7,000 frames available for linear regression analysis after downsampling in the cups and social assays; see below) were excluded; when necessary, assays were repeated on subsequent days.

#### Calcium imaging

Recordings of GCaMP6f activity were obtained at 20 Hz using Inscopix miniaturized microscopes and acquisition system (predominantly nVoke 2.0; 5 mFC-implanted mice were recorded using nVue 1.0. We switched to the nVoke 2.0 system for the majority of experiments due to improved signal quality and a higher yield of identifiable cells). Awake mice were connected to the recording system via commutators in passive mode, and the miniature microscope (Inscopix) was attached to the implanted baseplate prior to each experiment. For assays conducted in the empty arena (30 min), recording was continuous throughout the session. For the cups and social assay, neural activity was recorded during each 3-min session, paused during animal or cups repositioning between sessions, and then restarted, resulting in a total recording duration of 72 min for assays with cup or social landmarks. LED illumination was set to the lowest level that reliably captured calcium transients (typically 0.4-0.7, with most recordings *<* 0.7). The focal plane (Z-focus) was adjusted across sessions to maintain a consistent field of view, using anatomical landmarks such as blood vessels. Following each 72-min or 30-min experiment, the microscope was removed and replaced with a baseplate cover or dummy microscope before returning the mice to their home cages. Overhead video was acquired using a Blackfly S USB3 camera (1.6 MP, monochrome; Teledyne) and synchronized with calcium imaging via a hardware trigger from the Inscopix data acquisition (DAQ) system, which generated a pulse after capturing each imaging frame. Hardware-triggered acquisition and synchronization were implemented using custom Python code adapted from https://github.com/nicthib/FLIR-Multicam. Video was thus aligned frame-by-frame with the imaging data, and both were acquired at 20 Hz. Sessions in which synchronization failed were excluded from analysis. The mouse’s nose and tail-base were tracked using DeepLabCut, a pose-estimation software (42).

#### Imaging data processing

Motion correction and cell extraction were performed using a custom pipeline based on previously described methods (43), with minor modifications. Briefly, raw calcium imaging videos were spatially downsampled (4×) and processed to enhance stable anatomical features (e.g., blood vessels) using spatial band-pass filtering. This was followed by a difference-of-Gaussians filter to further emphasize high-contrast, stationary landmarks, as described previously (43). In brief, images were smoothed using a Gaussian filter (imgaussfilt, MATLAB) with a small kernel (radius 2 pixels for our downsampled 200×320 pixel images), and the result was subtracted from images smoothed with a broader kernel (radius 6 pixels) to emphasize high-frequency, stationary features. Pixel intensities were then thresholded (values *<* 4 set to 4). This yielded videos with a largely suppressed background and high-contrast landmarks, which were used for motion correction; the resulting shifts were then applied to the corresponding unfiltered videos. For each recording day, an anchor 3-min session was motion-corrected to generate a reference template using the NoRMCorre algorithm (44); this corresponded to a middle 3-min session for the cups and social assays and the full session for the 30-min empty arena assay. Motion correction was then performed on the filtered (difference-of-Gaussians) videos from all other sessions relative to this template, using NoRMCorre with the following parameters: bin_width = 500, init_batch = 1000, max_shift = 10, and iter = 2. The shifts were subsequently applied to the corresponding unfiltered (raw) videos. For both the social and cups assays, individual 3-min motion-corrected session videos were concatenated prior to further processing.

Motion-corrected videos were then processed using the MATLAB implementation of CNMF-E to extract spatial footprints and temporal activity of individual neurons (45). CNMF-E default parameters were used, except for min_corr = 0.8, min_pnr = 9, gSig = 3, and gSiz = 11. For initialization, min_pixel was set to 25. For identification of cells in the residual after background subtraction, min_corr_res and min_pnr_res were set to 0.8 and 9, respectively. Following two iterations of CNMF-E, additional filtering steps were applied to remove false positives as in (43), based on criteria implemented in the CaImAn Python package (46). Briefly, cells were filtered based on temporal sparsity (*>* 0.003), peak-to-noise ratio (*>* 8.5), and stability of baseline fluorescence across the session. Baseline stability was assessed by computing the difference in mean activity between the first and last 10% of session frames; cells with absolute differences greater than 3 were excluded. Putative neurons were further curated by visual inspection using the CNMF-E *viewNeurons* interface to confirm appropriate spatial footprints and calcium dynamics, and remaining false positives were removed.

CNMF-E performs frame-by-frame background subtraction and deconvolution, producing a denoised calcium trace (C) and an inferred spiking trace (S). For all analyses except spatial information content (Skaggs information) and determining the native decay constants of neurons, we used CNMF-E-extracted event amplitudes from S convolved with a fixed exponential kernel (2s decay constant), rather than raw fluorescence traces, and z-scored across the entire experimental session. This procedure standardizes decay kinetics across neurons and animals and enhances signals from temporally sparse activity. Hereafter, “z-scored calcium activity” refers to this 2-s kernel-convolved CNMF-E event trace after z-scoring across the full experimental session. Spatial information content was computed using inferred spike events as a mutual information measure between activity for each cell and the animal’s position (8, 47), because this measure assumes discrete, sparse activity and can be biased when applied to continuous fluorescence traces.

#### Estimation of the native decay time constant of each individual neuron

To estimate decay time constant (*τ*), we divided each neuron’s calcium activity (z-scored denoised calcium trace, C obtained from CNMF-E) into temporal segments bounded by consecutive inferred spike events (non-zero entries in S). Segments shorter than 2 s or lacking a clear decay (i.e., the z-scored signal remaining entirely above zero) were excluded. For each remaining segment, the signal was normalized to the range [0, 1] and fit with a single-exponential decay to estimate *τ*. This yielded a distribution of *τ* values per neuron, and the median *τ* was taken as the neuron’s characteristic *τ*.

#### Principal component analysis

We used principal component analysis (PCA) to examine the organization of spatial population activity. For each neuron, we computed an 8 × 8 spatial rate map by averaging z-scored calcium activity within each spatial bin. PCA was performed separately for individual animals and on neurons pooled across animals within each brain region and assay. Individual-animal PCA results were used for the decoder-PC alignment analysis (see below), whereas pooled-neuron PCA was used to characterize population-level spatial structure. Spatial maps were vectorized across bins and concatenated across neurons, such that spatial bins were observations and neurons were features. PCA was then performed by eigendecomposition of the neuron-by-neuron covariance matrix.

To quantify the dimensionality of population variance, we calculated the participation ratio of the full PCA variance spectrum for each animal:

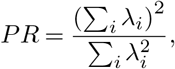

where *λ*_*i*_ is the fraction of total population variance explained by principal component *i*. Larger participation ratios indicate that population variance is distributed across a greater number of principal components, whereas smaller values indicate that variance is concentrated in fewer components. Participation ratios were compared across brain regions using a pairwise two-sided Welch’s *t*-test.

#### Alignment of linear position decoders with principal components

To determine how the spatial information used by linear position decoders was distributed across principal components, we measured the alignment between each decoder’s weight vector and the principal component loading vectors obtained from whole-arena PCA performed separately for each animal. For each animal, we used the linear decoders trained to predict *x* from the upper and lower halves of the arena, and *y* from the left and right halves. Decoder weight vectors were matched to the neurons included in PCA, preserving their original ordering.

For each decoder and principal component, alignment was quantified as the absolute cosine similarity between their weight vectors,

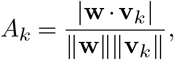

where w is the decoder weight vector and v_*k*_ is the loading vector of principal component *k*. Absolute cosine similarity was used because the sign of each principal component is arbitrary. We calculated alignment across principal components in order of decreasing explained variance.

To quantify how broadly decoder alignment was distributed across principal components, we calculated the participation ratio of the absolute cosine similarity values across the first 20 PCs:

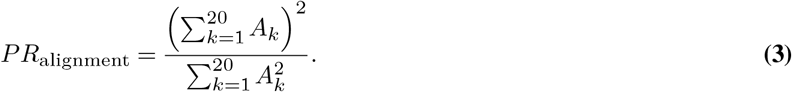

Lower values indicate that decoder alignment is concentrated in fewer principal components, whereas higher values indicate a broader distribution. Participation ratios were calculated separately for decoders trained in each half of the arena and then averaged within each animal for the *x* and *y* decoders.

To assess the consistency of decoder alignment across spatial regions, we calculated the Pearson correlation between the absolute cosine similarity profiles of decoders trained in opposite halves of the arena, using the first 60 PCs. Higher correlations indicate that decoders trained in different spatial regions rely on similar distributions of principal components. Each animal contributed one participation ratio and one alignment-profile correlation for each spatial axis. Differences between brain regions were assessed using pairwise two-sided Welch’s *t*-tests.

#### Singular value decomposition (SVD) and quadratic fitting of population spatial activity maps

To characterize the dominant spatial patterns of neuronal activity, we pooled the spatial activity maps of all neurons across animals within each brain in mFC and dCA1, separately for the cups and social conditions. Each neuron’s activity map was represented on an 8 × 8 spatial grid. The four corner bins were excluded, leaving 60 spatial bins per neuron.

We constructed a 60 × *N* matrix, where *N* is the total number of neurons pooled across animals, and each column corresponds to the spatial activity map of one neuron. The matrix was mean-centered by subtracting the mean activity across neurons at each spatial bin. We then performed singular value decomposition (SVD) on the centered matrix. The first three spatial modes were obtained by multiplying the corresponding left singular vectors by their singular values.

To assess whether these spatial modes could be described by quadratic functions of position, we fitted each mode independently using the model

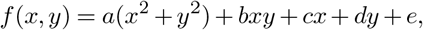

where *x* and *y* are spatial coordinates normalized to the range [−1, 1], and *a, b, c, d*, and *e* are fitted coefficients. The coefficients were estimated by ordinary least squares across the 60 spatial bins. RMSE threshold was quantified using the coefficient of determination (*R*^2^) between each spatial mode and its fitted quadratic function. To visualize the contributions of the individual spatial terms, we separately plotted *a*(*x*^2^ + *y*^2^), *bxy, cx*, and *dy* for each fitted mode.

#### Reconstruction of spatial functions from the first three SVD modes

To determine whether the first three spatial modes could jointly represent the spatial functions predicted by the squared-distance model, we used ordinary least squares regression to reconstruct *x, y*, and *x*^2^ + *y*^2^ from linear combinations of these modes. Each spatial function was fitted separately using

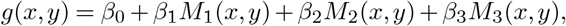

where *g*(*x, y*) denotes *x, y*, or *x*^2^ + *y*^2^, and *M*_1_, *M*_2_, and *M*_3_ are the first three singular-value-weighted spatial modes. The coefficients were estimated by ordinary least squares across the 60 spatial bins, using the same normalized coordinates as in the quadratic fitting analysis. Reconstruction accuracy was quantified using the coefficient of determination (*R*^2^) between each target function and its fitted reconstruction. For visualization, we plotted the reconstructed spatial maps without the fitted intercept.

#### Spatial information

Spatial information content was computed from inferred spike events using the Skaggs information measure. For each cell, the arena was divided into an 8 × 8 spatial grid, and a spatial rate map was constructed by dividing the number of inferred events in each spatial bin by the total number of frames spent in that bin. Here, inferred events were defined by thresholding the CNMF-E-derived deconvolved spike trace (S) to obtain a binary event vector. Spatial information was then calculated as:

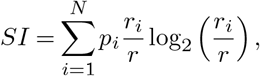

where *p*_*i*_ is the occupancy probability of spatial bin *i, r*_*i*_ is the mean event rate in that bin, and *r* is the overall mean event rate across the experiment.

To assess significance, inferred event trains were circularly shifted relative to the animal’s trajectory 1000 times for each cell, and spatial information was recomputed for each shuffle. Cells were classified as spatially selective if the observed spatial information exceeded the shuffle distribution by a z-score greater than 3.

#### Linear regression decoding of spatial position

All decoding and generalization analyses were performed separately for each animal.

For each animal, z-scored calcium activity was represented as an N x T matrix, where N is the number of neurons and T the number of time points. We then trained linear regression models to predict the animal’s frame-by-frame nose position along either the *x*- or *y*-axis from population activity vectors. X and Y axes were defined relative to a fixed arena reference frame, with the door located on the left side of the arena, such that *x*-values increase from left to right and *y* values increased from top to bottom.

For full-arena decoding (Fig. 1d-g), models were trained and evaluated using frames spanning the entire arena. Decoding was performed independently for *x*- and *y*-position. To control for non-uniform spatial sampling, frames were balanced with respect to the decoded coordinate: for *x*-decoding, the arena was divided into four vertical strips spanning the *x*-axis, and an equal number of frames was randomly sampled from each strip; for *y*-decoding, the arena was divided into four horizontal strips spanning the *y*-axis and sampled analogously. Decoding performance was then evaluated using 10-fold cross-validation with contiguous folds (KFold, shuffle=False). The data were divided into 10 consecutive temporal blocks; on each fold, the model was trained on nine blocks and tested on the remaining block, with each block serving once as the test set. Predictions from held-out folds were concatenated to obtain predicted positions across the session. Analyses were restricted to moving frames, defined using centroid speed *>* 2 cm/s, with centroid position taken as the midpoint between nose and tail-base coordinates. Decoding performance was quantified as the median absolute error between true and predicted position and converted from pixels to centimeters using a calibration factor of 14.49 pixels/cm (constant across all videos due to the fixed camera position relative to the arena). As a null model, neural activity for each neuron was independently circularly time-shifted, preserving its temporal autocorrelation and activity distribution while disrupting its alignment to behavior. 100 shuffled datasets were generated for each animal and analyzed using the same procedure. For all decoding and generalization analyses, observed performance was compared across animals with each animal’s mean shuffle performance using two-sided paired *t*-tests.

For generalization decoding of *x*-position, we trained on frames when the nose was on the top/bottom of the arena and tested on frames when the nose was on the other half, ensuring that *x* position could be predicted independently of *y*. Conversely, for generalization decoding of *y*-position, we trained on frames when the nose was on the left or right half and tested on frames when the nose was on the other half, ensuring that *y* could be predicted independently of *x*. This partitioning was critical for evaluating generalization, as it ensured that the coordinate to be decoded spanned its full range within each half, while the orthogonal coordinate was restricted, enabling assessment of generalization to spatial regions that were out of distribution with respect to the orthogonal coordinate. Spatial partitions were further defined with a 5 percent gap between adjacent halves to exclude frames near the dividing boundary. In addition, brief crossings between adjacent halves were removed: if the animal traversed the gap in less than 2 s (40 frames at 20 Hz), frames surrounding the transition were symmetrically excluded to yield a total exclusion window of 2 s. To control for non-uniform sampling of the decoded coordinate, frames were spatially balanced prior to decoding, similar to full-arena decoding. For *x*-position, frames from the top and bottom halves were separately subdivided into four vertical strips spanning the *x*-axis, and equal numbers of frames were randomly sampled from each strip. For *y*-position, frames from the left and right halves were analogously subdivided into four horizontal strips spanning the *y*-axis and equally sampled across *y*-ranges. Following downsampling, experiments with fewer than 7,000 frames available for either coordinate were excluded. Generalization was evaluated in both directions: for *x*-position, models were trained on the top half and tested on the bottom half, and vice versa; for *y*-position, models were trained on the left half and tested on the right half, and vice versa. Generalization performance was quantified on the test set as the median absolute error between true and predicted position. Median generalization errors were computed for each train/test configuration, and their mean is reported as generalization errors in the figures. Decoding within individual spatial partitions was evaluated using the same 10-fold contiguous cross-validation procedure described above, separately for each half and each coordinate. The median error was calculated within each half, and the mean across the two halves was reported. The same movement restriction, spatial downsampling procedure used to balance sampling along the decoded coordinate, exclusion criteria, and circular time-shift shuffle procedure described above for full-arena decoding were applied to both within-partition decoding and cross-partition generalization. For comparisons across experimental conditions (empty, cups, and social), analyses were restricted to the first 30 min of each session.

#### Weight consistency across arena halves

For each recording session, separate linear models were trained on the two halves of the arena for decoding*x*- and *y*-position. For each spatial axis, decoder weights from the two arena halves were normalized jointly by the Euclidean norm of the concatenated weight vectors. Neurons were then pooled across animals within each brain region. Decoder-weight consistency across halves was quantified as the Pearson correlation (*r*) between the corresponding population weight vectors, separately for each spatial axis.

To assess whether neurons well described by the squared distance model contributed more consistently to position decoding, we divided neurons into two groups based on the model’s RMSE threshold. For this analysis, each decoder’s weight vector was normalized independently by its Euclidean norm. Within each group, we computed the Pearson correlation between decoder weights obtained from opposite halves of the arena, separately for *x*- and *y*-position.

To examine whether differences in calcium dynamics could account for decoder-weight consistency across arena halves, we tested the relationship between each neuron’s native calcium decay constant and the consistency of its decoder weights across halves. We found no strong correlation between these measures (Extended Data Fig. 13).

#### Bayesian decoding of two-dimensional position

A Bayesian decoder was used to decode the animal’s two-dimensional position from z-scored calcium activity, as described above. Neural activity and position were averaged in non-overlapping 250-ms bins, and bins were included when at least 80% of frames were classified as movement. The arena was divided into 8 × 8 spatial bins. For each neuron, mean activity was estimated separately for each spatial bin, whereas a single activity variance was estimated across all training samples. Spatial rate maps were smoothed with a 2D Gaussian kernel (*σ* = 1; kernel truncated at 3*σ*), and spatial bins containing fewer than five training samples were excluded from decoding.

The decoder assumed Gaussian activity distributions and conditional independence between neurons. For each temporal bin, the posterior probability of the animal occupying spatial bin *s* was computed as

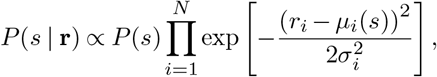

where r is the population activity vector, *r*_*i*_ is the activity of neuron *i, µ*_*i*_(*s*) is its mean activity in spatial bin *s*, 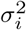 is its activity variance, and *P* (*s*) is the spatial prior. A uniform prior was used across valid spatial bins. The center of the bin with the maximum posterior probability was selected as the decoded position.

Decoder performance was evaluated using five-fold cross-validation, with the recording divided into five contiguous temporal blocks. Four blocks were used for training and the remaining block for testing, with each block serving once as the test set. Decoding error was calculated as the Euclidean distance between the true and decoded positions. For each cross-validation fold, the mean decoding error was calculated across all held-out samples. For each animal, decoding performance was summarized as the mean of the five fold-wise mean decoding errors. To generate a null distribution, neural activity was circularly shifted relative to behavior 100 times, with shifts of at least 30 s. For each brain region and assay, observed decoding errors were compared with the corresponding mean shuffled decoding errors across animals using a two-sided paired *t*-test, with each animal contributing one observed value and the mean of its 100 shuffled values. Differences in decoding error across brain regions were assessed using one-way ANOVA followed by pairwise two-sided Welch’s *t*-tests with Holm correction for multiple comparisons.

#### From generalized quadratic distances to *x* and *y* coordinates

Here we show that, for the generalized quadratic distance model of neural responses, it is possible to linearly read out the animal’s *x* and *y* coordinates by computing a linear combination of neuronal activities.

More formally, we consider a population where neurons are grouped into *K* distinct landmarks. Let (*x*_*k*_, *y*_*k*_) denote the coordinates of the *k*-th distinct landmark, and let *f*_*k*_(*x, y*) be the response function for neurons tuned to this landmark, given by:

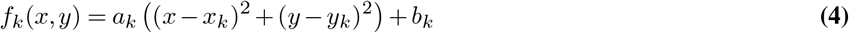

Expanding this expression yields:

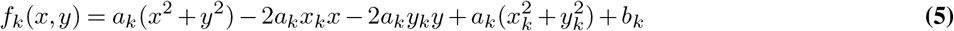

We seek to determine the geometric and parametric conditions under which a downstream linear readout can decode the spatial coordinates *x* and *y*. For the *x*-coordinate, we require a set of effective synaptic weights *W*_*k*_ for each landmark such that:

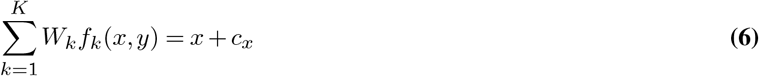

where 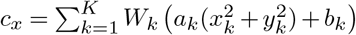 is a constant bias term.

To isolate the *x* term and eliminate both the nonlinear quadratic term and the orthogonal *y* term, substituting the expanded form of *f*_*k*_(*x, y*) and matching coefficients requires the weights to satisfy the following three constraints:

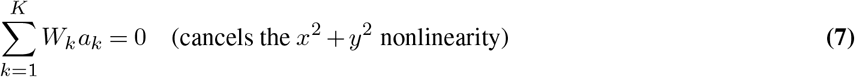

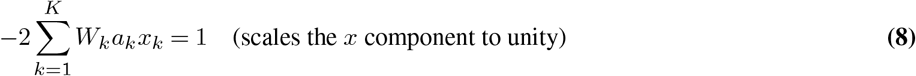

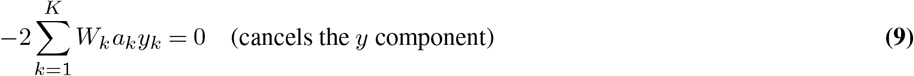

Rewriting these conditions in terms of a matrix equation **MW** = **b**_*x*_ yields:

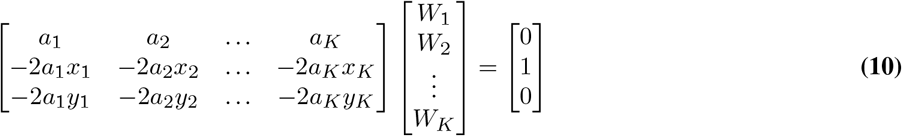

An analogous system, MV = b_*y*_ with target vector b_*y*_ = [0 0 1]^⊤^, must be solved to decode the *y*-coordinate.

##### Geometric and Parametric Conditions for Decoding

For these systems to admit a consistent solution across arbitrary spatial trajectories, the constraint matrix M ∈ ℝ^3×*K*^ must have full row rank, i.e., rank(M) = 3. This imposes requirements on both the spatial arrangement and the scaling parameters (*a*_*k*_):

1. **Dimensionality (***K* ≥ 3): We require at least *K* ≥ 3 distinct landmarks with non-zero scaling (*a*_*k*_ ≠ 0). If *K <* 3, the column space of M is at most two-dimensional, preventing general alignment with the target readout vectors.
2. **Non-collinearity:** The landmarks cannot all lie on a single straight line. If *y*_*k*_ = *mx*_*k*_ + *c* for all *k*, the third row of M remains a linear combination of the first two rows (*R*_3_ = *mR*_2_ + *cR*_1_) regardless of the *a*_*k*_ multipliers. This reduces the rank of M to at most 2. Thus, at least three landmarks with independent spatial locations must form a non-degenerate triangle.
3. **Parametric Heterogeneity:** The scaling factors *a*_*k*_ modulate the effective weight contributions in each row. If all *a*_*k*_ are identical (e.g., *a*_*k*_ = *a*), they factor out as scaling constants on the rows, preserving the rank structure of the original unscaled model. However, if *a*_*k*_ vary across landmarks, they introduce additional weighting that can affect conditioning, though the rank condition is still governed primarily by the spatial coordinates (*x*_*k*_, *y*_*k*_) as long as *a*_*k*_ ≠ 0.

#### Singular Value Decomposition of populations of quadratic distance neurons

Under the assumption that all neurons are encoding the quadratic distance from landmarks, we can show that the data matrix can be easily decomposed in a way that clearly reveals: 1) that the dimensionality is always 3; 2) each component is a linear superposition of the *x, y*, and *x*^2^ + *y*^2^ terms.

Consider a population of *N* neurons where each neuron *k* encodes the quadratic distance from a specific landmark located at coordinates (*x*_*k*_, *y*_*k*_). The firing rate of neuron *k* as a function of the animal’s position (*x, y*) is given by:

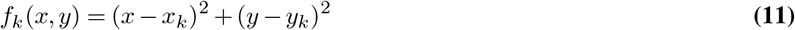

For simplicity, we assume that the *a*_*k*_ coefficient of the model defined in Fig 3 is 1, and *b*_*k*_ = 0. Expanding this expression yields:

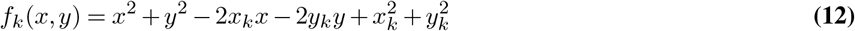

We assume that the arena is discretized and that the animal uniformly visits the *M* discrete spatial bins (indexed by *m* = 1,…, *M*). The bins have spatial coordinates *x*(*m*) and *y*(*m*). The *f*_*k*_(*x*(*m*), *y*(*m*)) for all *k* and *m* form the data matrix *X* (of dimensions *N* × *M*, where rows represent neurons and columns represent spatial bins).

Standard Principal Component Analysis (PCA) and Singular Value Decomposition (SVD) require mean-centering each neuron’s activity across space. We subtract from every *f*_*k*_ the average activity across all spatial bins, obtaining a new centered *f*_*k*_ given by:

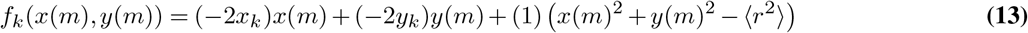

where ⟨*r*^2^⟩ is the averaged squared radial distance across the arena. Each neuron’s centered field is strictly confined to a 3-dimensional subspace spanned by the spatial basis functions evaluated across the arena: 1) The linear *x*-coordinate: *x*(*m*) 2) The linear *y*-coordinate: *y*(*m*) 3) The mean-centered radial squared term: *x*(*m*)^2^ + *y*(*m*)^2^ −⟨*r*^2^⟩

##### Matrix Factorization and Spatial Covariance

The population data matrix *X* can be formulated as a matrix product:

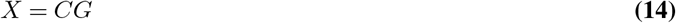

where *G* is the 3 × *M* spatial basis matrix, containing the evaluated basis functions across all *M* spatial bins (*m* = 1,…, *M*):

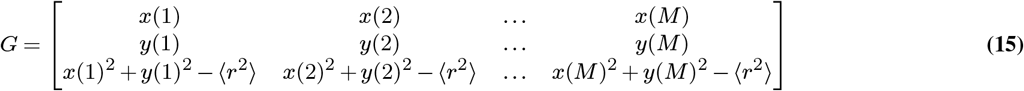

and *C* is the *N* × 3 neural weight (coefficient) matrix, containing the landmark-dependent parameters for all *N* neurons (*k* = 1,…, *N*):

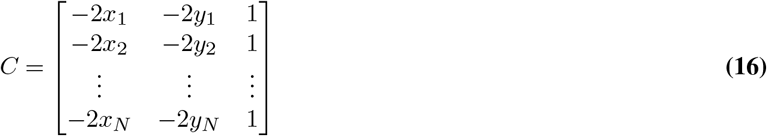

Notice that *C* = *M*^*T*^, where *M* is the matrix introduced and discussed in the previous section. This decomposition already shows that the dimensionality is 3, both in the neural activity space (in which each point represents one position, and the coordinate axes represent the activities of different neurons) and in the fields space (in which each point is one neuron and the coordinate axes represent the activities at different locations). Indeed, the rank of the product of two matrices is always smaller than or equal to the minimum of the ranks of the two matrices, which is 3, assuming that *N* and *M* are large, as in the data.

##### Singular Value Decomposition (SVD) and Right Eigenvectors

To perform SVD on the data matrix *X* = *USV*^*T*^, we analyze the spatial covariance matrix *X*^*T*^ *X* (of size *M* × *M*):

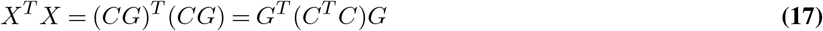

The right singular vectors (columns of *V*, living in the *M*-dimensional spatial bin space) form an orthonormal basis for the spatial tuning profiles. Their exact form depends on the distribution of landmarks:

##### Uniform Arena and Symmetric Landmarks

If the arena is a symmetric square [−*L, L*] ×[−*L, L*] and the landmark coordinates {(*x*_*k*_, *y*_*k*_)} are distributed symmetrically such that cross-terms vanish 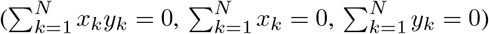, both *GG*^*T*^ and *C*^*T*^ *C* are diagonal matrices.

Consequently, the right singular vectors directly recover the orthogonal spatial basis functions: two degenerate singular vectors corresponding to pure linear gradients along *x* and *y* (due to isotropic variance where ⟨*x*^2^⟩_space_ = ⟨*y*^2^⟩_space_). A distinct third singular vector corresponding to the concentric radial quadratic map *x*^2^ + *y*^2^.

##### Arbitrary or Asymmetric Landmark Distributions

If the landmark positions are completely arbitrary or clustered, the neural weight covariance matrix *C*^*T*^ *C* becomes non-diagonal. While the row space of *X* remains strictly bounded within the same 3D subspace, the SVD will yield right singular vectors that are rotated or skewed linear mixtures of *x, y*, and *x*^2^ + *y*^2^. Notice that in this case, linear regression, for example, for *x*, will use a direction that is orthogonal to the *y* and *x*^2^ + *y*^2^ directions, and partially aligned to the *x* direction. When the three directions are linearly independent in the firing rate space (see under what conditions in the previous section), then this is always possible using Gram-Schmidt orthogonalization.

#### Single neuron analysis of the fields

Neural spatial representations were analyzed by modeling the firing rate of individual neurons as a function of the animal’s spatial coordinates (*x, y*). Raw empirical spatial firing rate maps, *Z*(*x, y*), and their associated standard deviation maps, *S*(*x, y*), were extracted for each neuron. The four unexplored extremities (corners) of the square arena were excluded from the fitting procedure. Spatial coordinates were normalized to the range [−1, 1] across both axes.

We fitted a pure distance model to the measured fields:

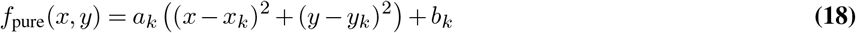

Model parameters were estimated using a weighted non-linear least squares optimization (trust-region-reflective algorithm). The residual objective function was weighted by the inverse of the empirical standard deviation at each spatial bin: *W* (*x, y*) = 1*/* max(*S*(*x, y*), *ϵ*). Initial parameter guesses were derived from a closed-form weighted linear least squares fit on the expanded polynomial forms to prevent convergence on local minima.

A fit was classified as valid (“well-fit”) if the Root Mean Square Error (RMSE) of the residuals—adjusted for the model’s degrees of freedom—fell below a threshold (75, 80, 85%) of the standard deviation of the spatial bins in the raw rate map (RMSE *<* 0.75, 0.8, 0.85*σ*_*Z*_).

## Acknowledgements

We thank Evan Schaffer and Jennifer Bussell for suggestions on motion correction, and members of the Axel lab for helpful discussions. We are grateful to Molly Rose Graff for assistance with histology, and to Nataliya Zabello and the staff of the Zuckerman Institute for Comparative Medicine for expert animal care. We further acknowledge Tanya Tabachnik, Mira Vanchiswar, and the Zuckerman Institute Advanced Instrumentation Core for assistance with modifications to the behavioral chambers for calcium imaging experiments, and the Zuckerman Institute’s Cellular Imaging platform for instrument use and technical advice. S.F. is grateful to Tim Behrens, Tom Mrsic-Flogel and their groups for many useful discussions during his visit to the Sainsbury Wellcome Center in London.

## Author Contributions

R.S., R.A. and S.F. conceived the project. R.S., A.M. and B.Z. performed the experiments. R.S., L.P., A.M., L.A., R.A. and S.F. analyzed the data. L.A., R.A. and S.F. supervised the project. R.S. wrote the initial draft of the manuscript. R.S., L.P., A.M., L.A., R.A. and S.F. contributed to writing and editing the manuscript.

## Extended Data

**Extended Data Fig. 1.**
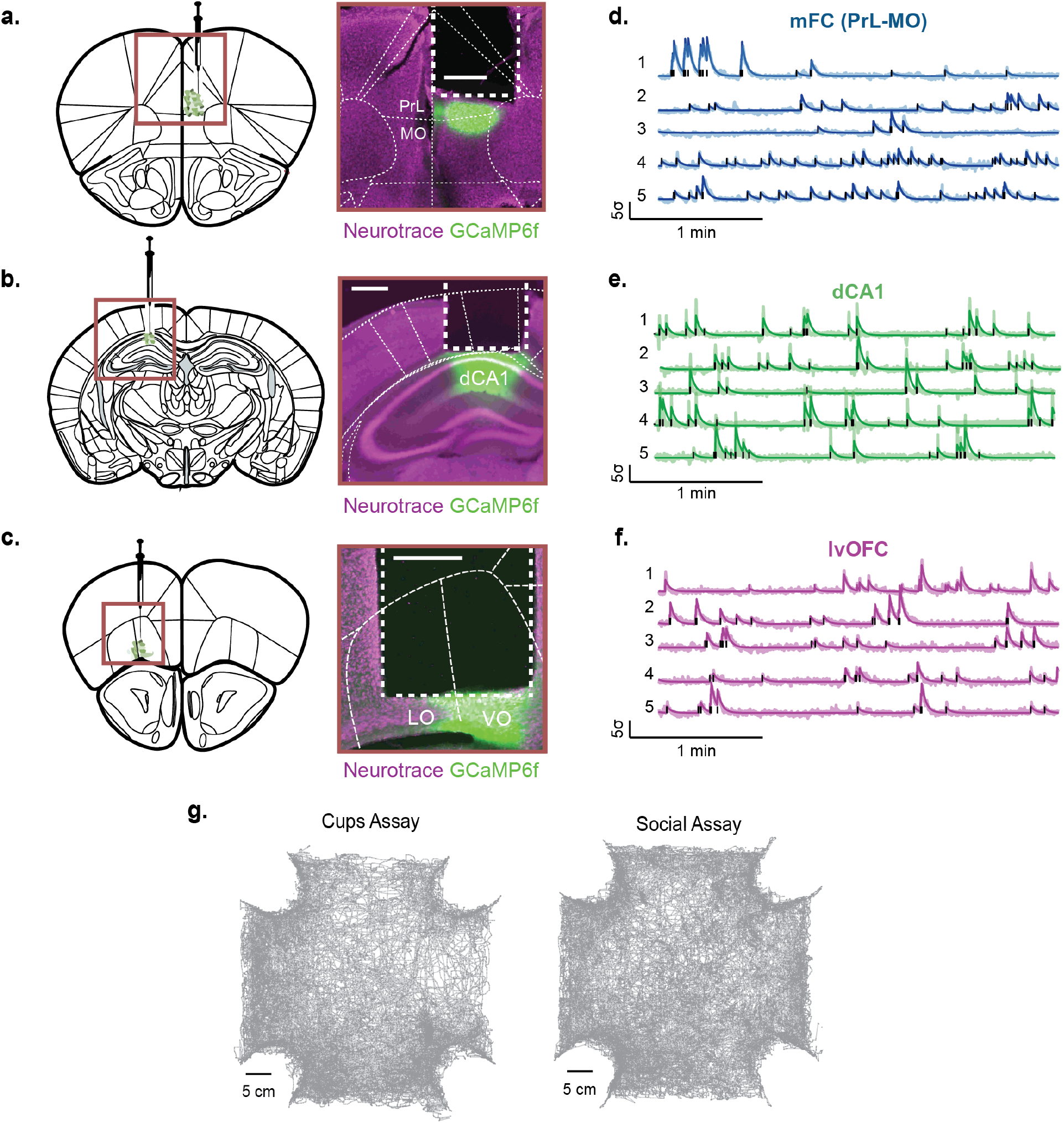
Recording sites and example behavioral trajectories. **a-c**. Coronal atlas schematics (left) and representative histological sections (right) showing GRIN lens placement and GCaMP6f expression in the prelimbic (PrL) and medial orbitofrontal (MO) regions of the medial frontal cortex (mFC; **a**), dorsal hippocampal CA1 (dCA1; **b**) and lateral and ventral orbitofrontal cortex (LO & VO) **c**). Sections were stained with Neurotrace (magenta) and show GCaMP6f fluorescence (green). Scale bars, 500 µm. **d-f**. Representative calcium activity traces from individual neurons recorded from mFC **(d)**, dCA1 **(e)**, and lvOFC **(f)** during exploration. Each row corresponds to a different neuron. Vertical black ticks indicate inferred calcium events obtained using constrained non-negative matrix factorization (CNMF-E). For all analyses, event amplitudes were convolved with a fixed exponential kernel (2-s decay constant; dark traces) rather than using raw fluorescence traces (light traces). This procedure standardizes decay kinetics across neurons and animals and enhances signals from temporally sparse activity. Scale bars, 1 s.d. (vertical) and 1 min (horizontal). **g**. Example nose position trajectories of a mouse over the full 72-min session in the cups assay (left) and social assay (right), comprising 24 consecutive 3-min sessions. Scale bar, 5 cm.

**Extended Data Fig. 2.**
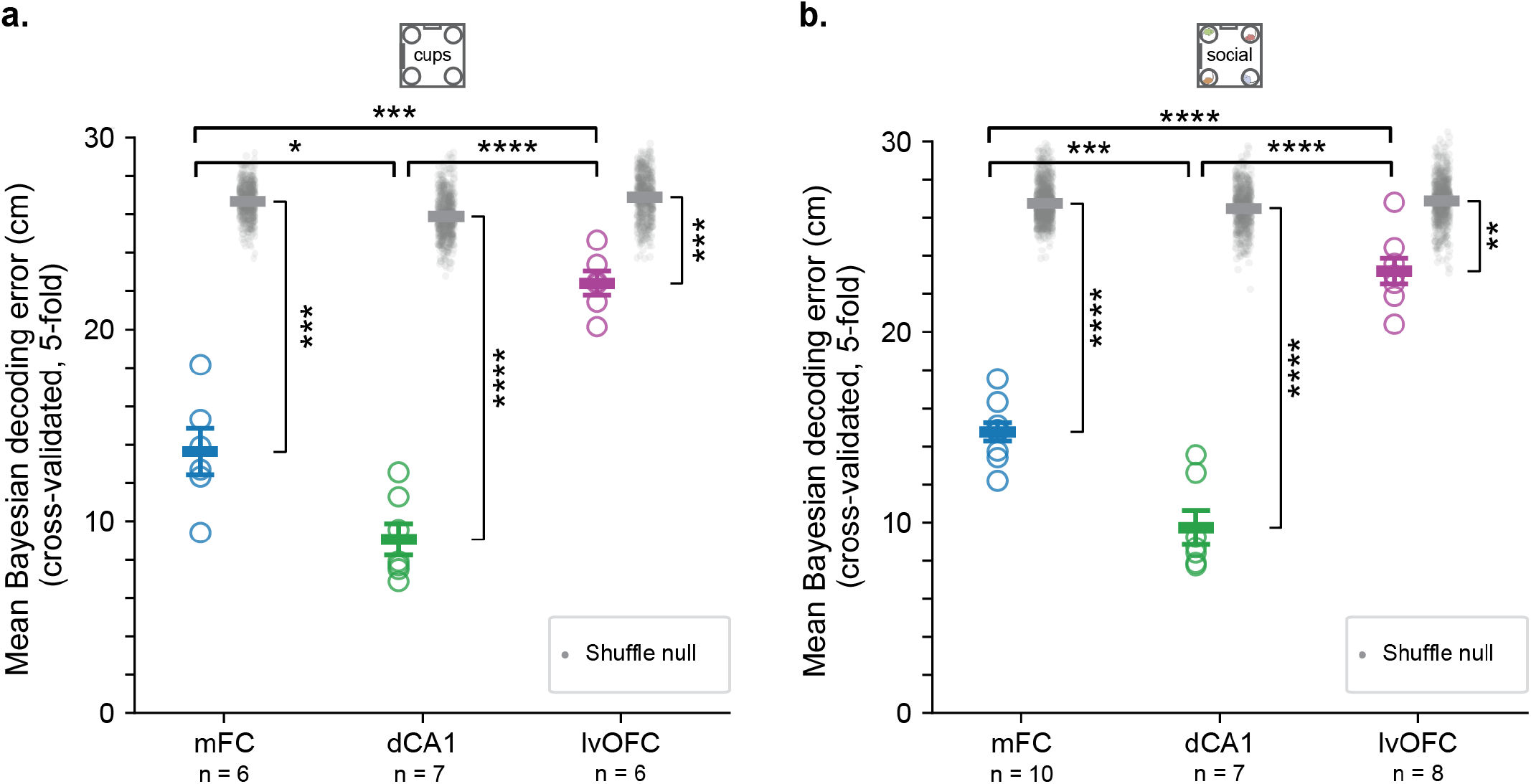
Bayesian decoding of position across brain regions. **a-b**. Mean cross-validated Bayesian position-decoding error for mFC, dCA1 and lvOFC recordings in the cups assay **(a)** and social assay **(b)**. Position was decoded using a two-dimensional Gaussian Bayesian decoder with five contiguous cross-validation folds, and decoding error was quantified as the mean Euclidean distance between the true and decoded positions. Colored circles indicate data from individual mice; horizontal lines and error bars indicate the mean ± SEM.

**Extended Data Fig. 3.**
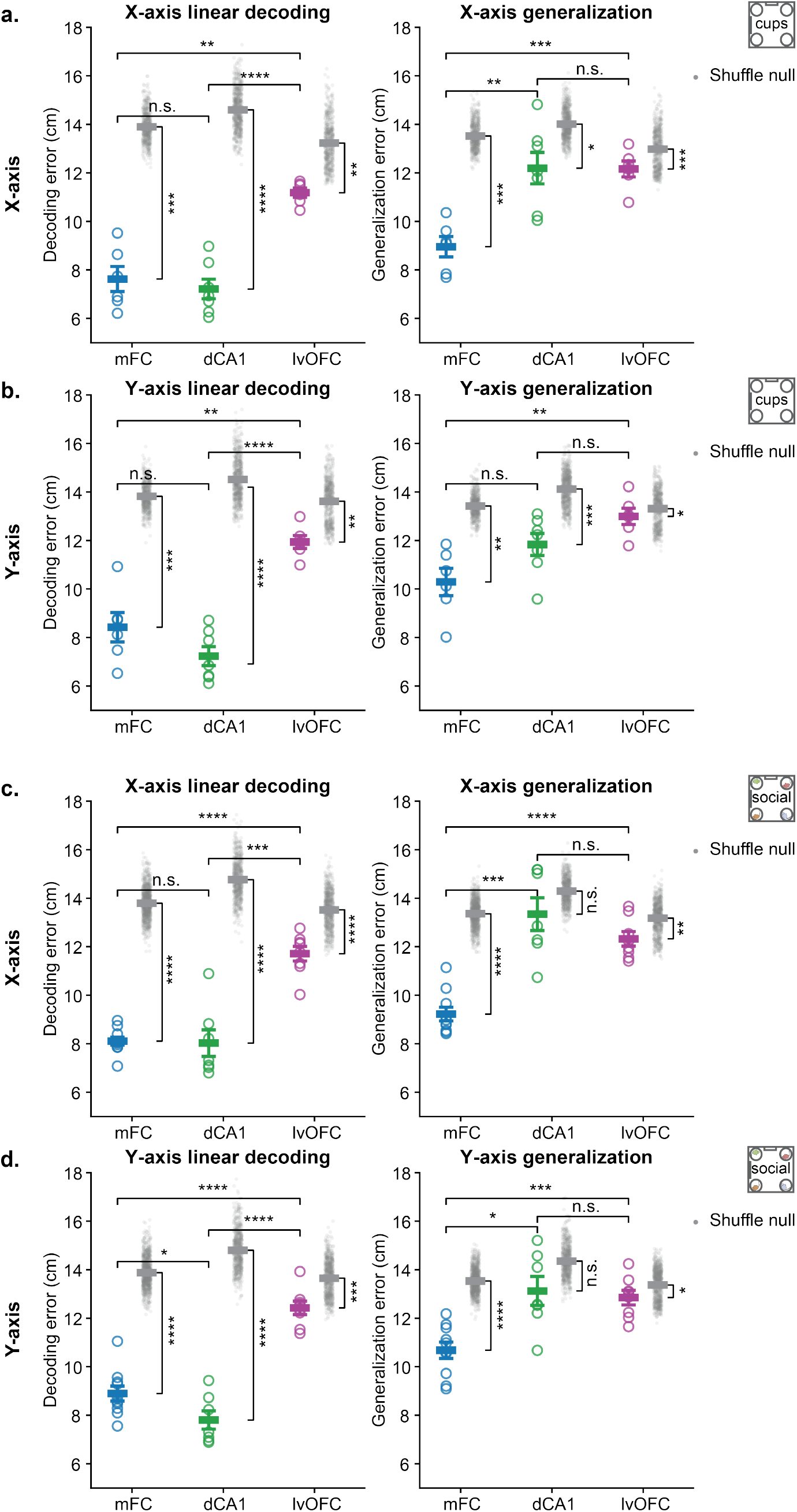
Linear decoding and cross-half generalization across brain regions. **a-d**. Comparison of linear decoding error (left), generalization error (middle) and the ratio of generalization to decoding error (right) across mFC, dCA1 and lvOFC for *x*-position in the cups assay **(a)**, *y*-position in the cups assay **(b)**, *x*-position in the social assay **(c)** and *y*-position in the social assay **(d)**. Each colored circle represents data from one animal; horizontal lines and error bars indicate the mean ± SEM. Decoding and generalization errors are quantified as in (Fig.2c-d) and replotted.

**Extended Data Fig. 4.**
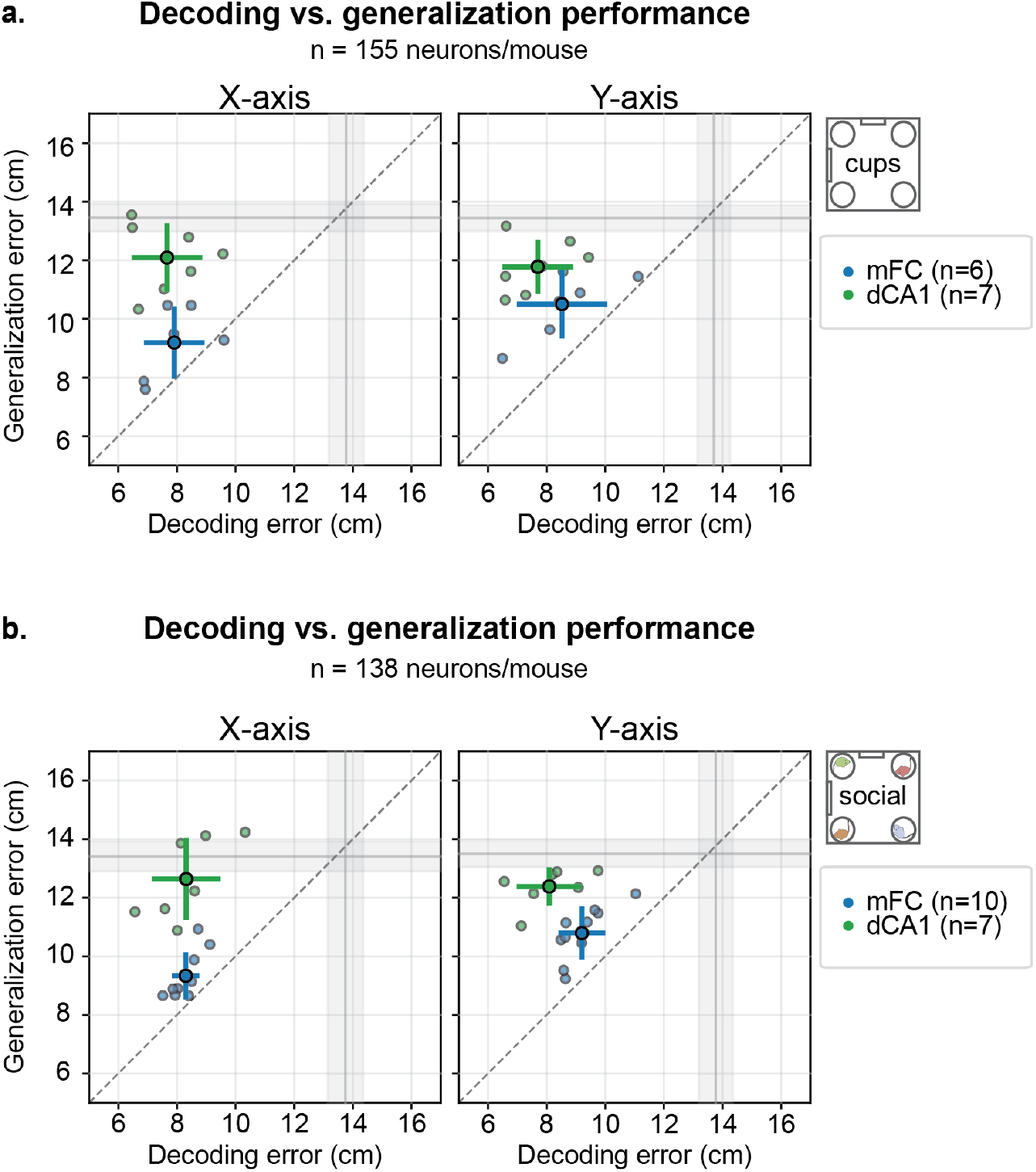
Decoding and generalization after matching population size across brain regions. **a,b**. Decoding versus generalization performance in the cups assay (**a**) and social assay (**b**) after matching population size across mFC and dCA1. For each assay, dCA1 neural populations were restricted to the minimum number of neurons recorded in an mFC animal (155 neurons per mouse for cups; 138 neurons per mouse for social) by selecting the top *N* neurons ranked by signal-to-noise ratio. Linear regressors were trained to predict horizontal (*x*; left) or vertical (*y*; right) nose position and tested either within the same arena half (decoding) or across opposite arena halves (generalization). Each dot represents one animal; larger dots and error bars indicate the mean ± SEM. Gray shading denotes the mean ± s.d. of shuffle-derived errors.

**Extended Data Fig. 5.**
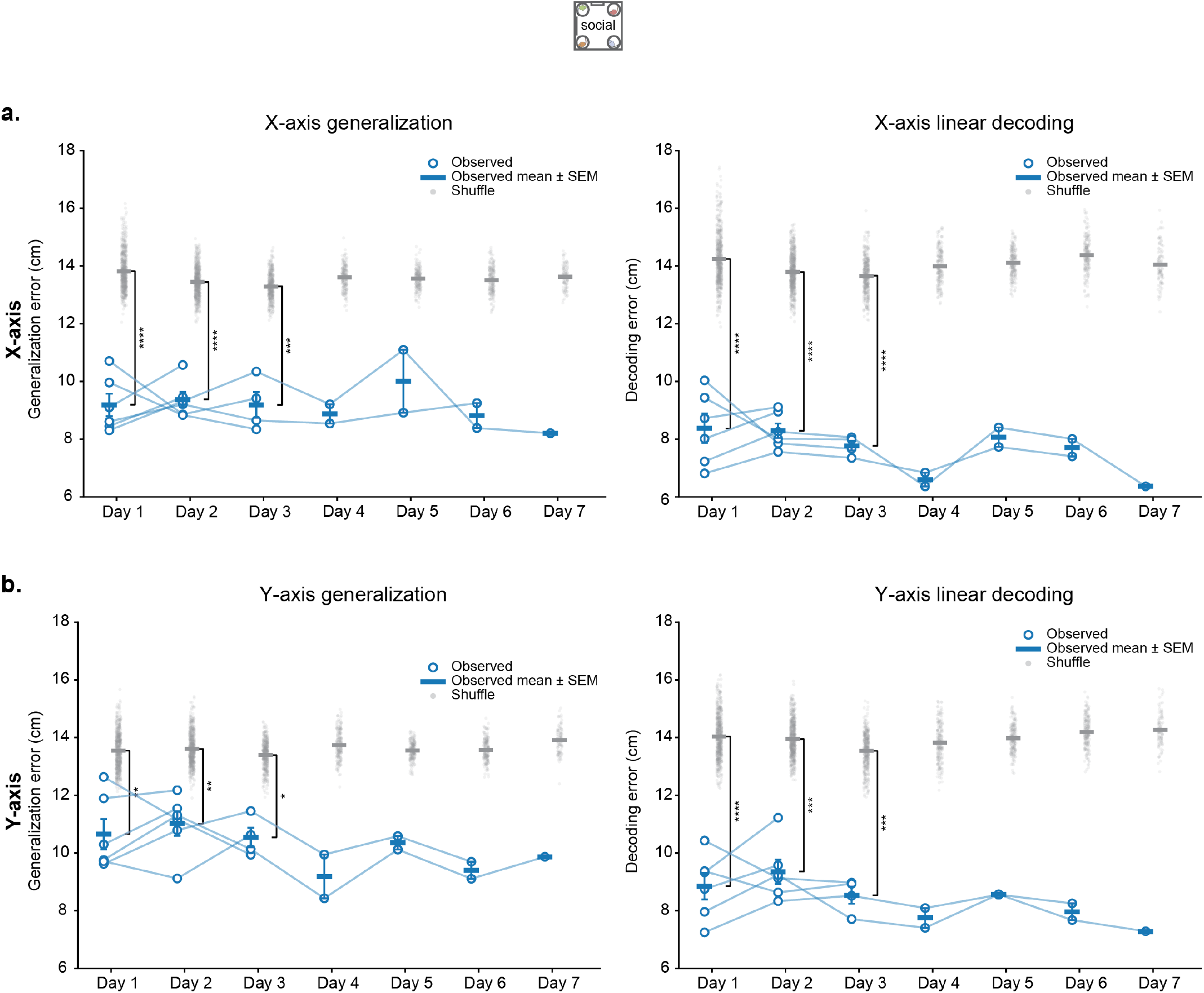
Stability of linear decoding and generalization across recording days in mFC. **a**. *x*-position generalization error (left) and within-half decoding error (right) across repeated recording days in the social assay. Open circles indicate individual mice, with lines connecting repeated measurements from the same mouse; horizontal bars and error bars indicate the mean ± SEM across mice. **b**. Same as **a**, for *y*-position generalization (left) and decoding (right), with generalization measured across left and right arena halves.

**Extended Data Fig. 6.**
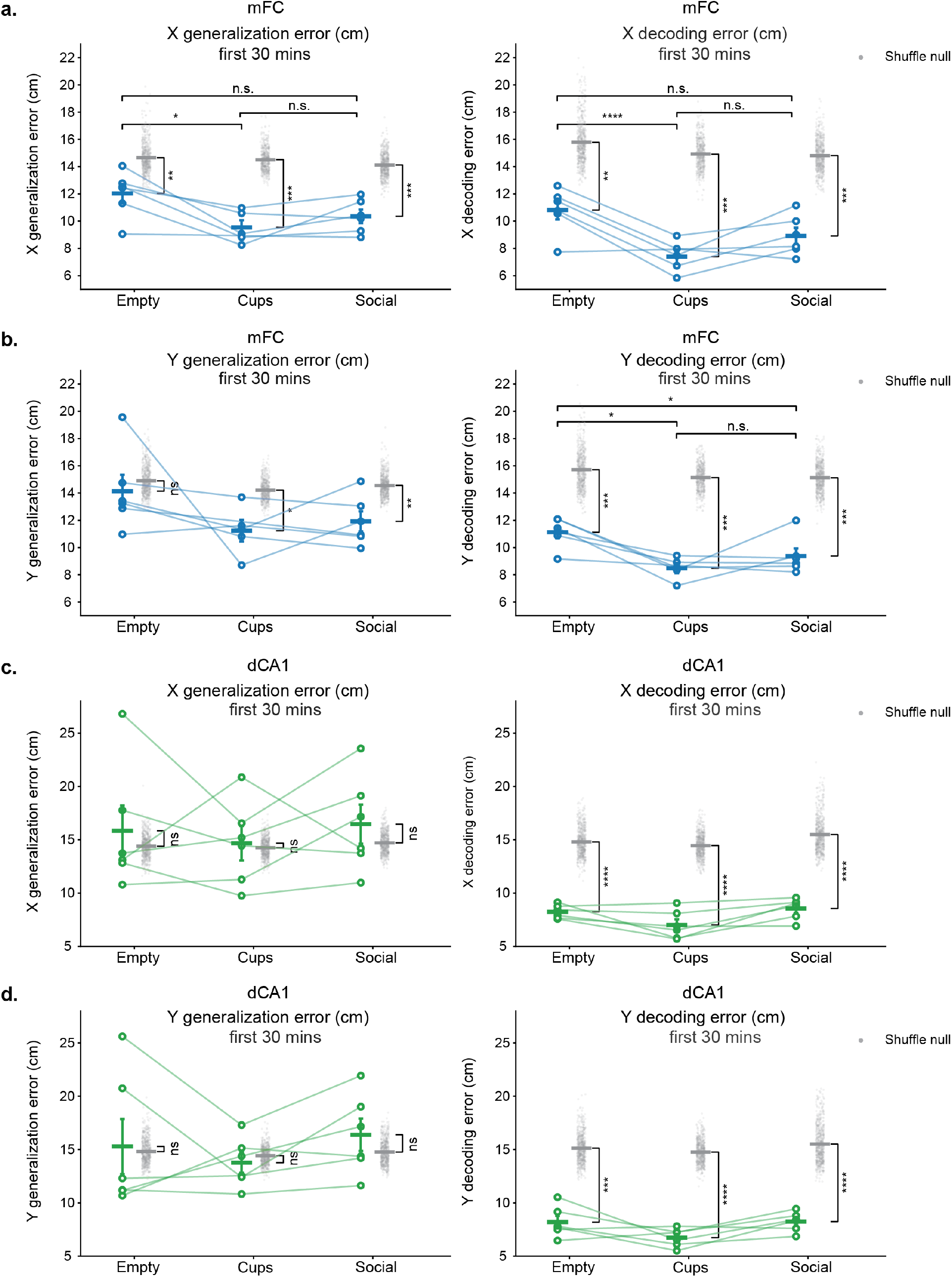
Linear decoding and generalization across assays in mFC and dCA1. **a**. *x*-position generalization error (left) and within-half decoding error (right) in mFC during the first 30 min of the empty-arena, cups, and social assays. Open circles indicate individual mice, with lines connecting measurements from the same mouse across conditions; horizontal bars and error bars indicate the mean ± SEM. **b**. Same as **a**, for *y*-position generalization error (left) and decoding error (right), with generalization measured across left and right arena halves. **c**,**d**. Same as **a**,**b**, respectively, for dCA1.

**Extended Data Fig. 7.**
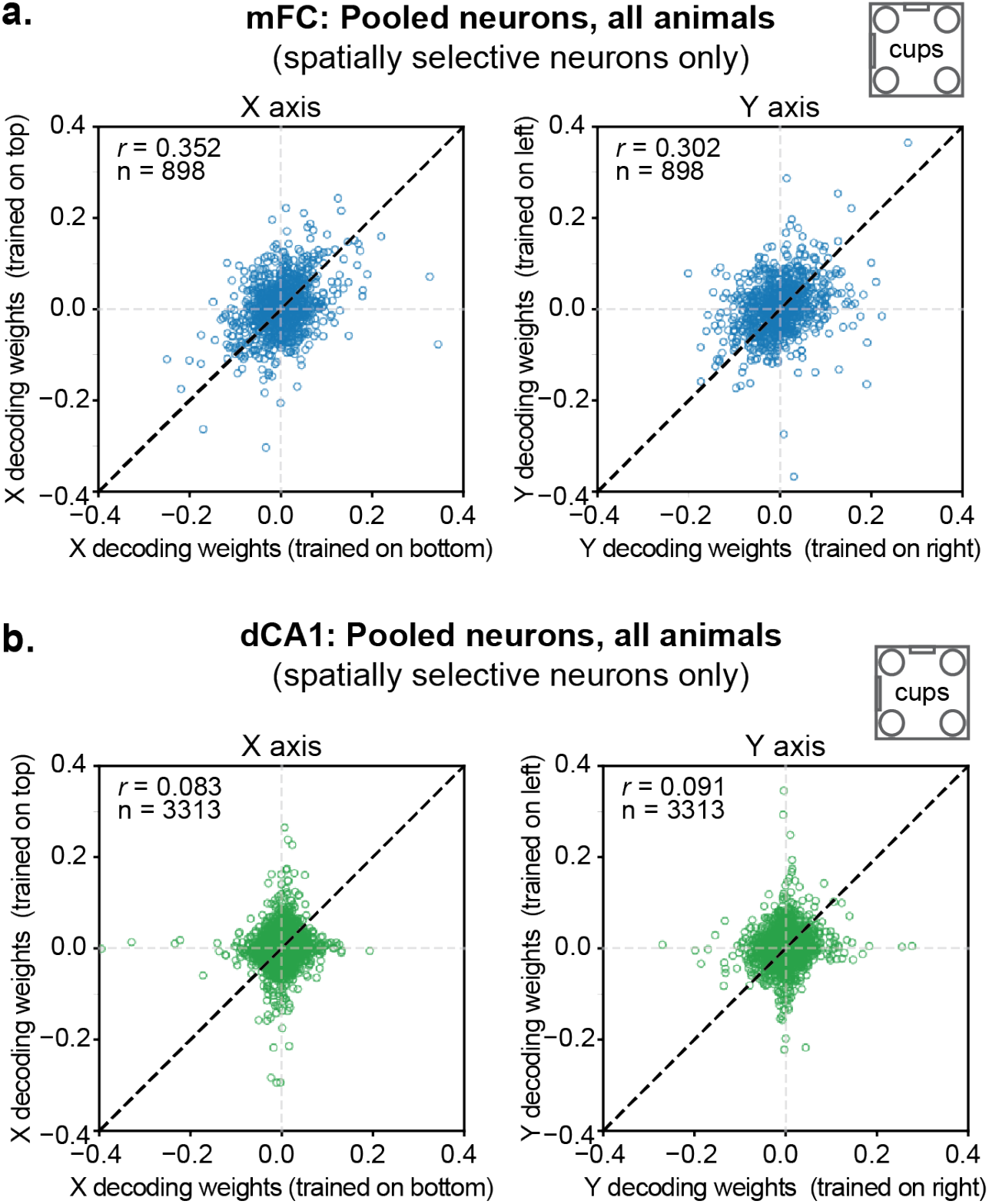
Cross-half weights analysis in the cups assay. **a-b**. Weights assigned to individual neurons by decoders trained on opposite halves are compared for *x*- (left) and *y*- (right) position for mFC (**a**) and dCA1 (**b**) populations in the cups assay. Neurons were pooled across animals. For each animal and spatial axis, the two decoder weights for each neuron were divided by the Euclidean norm of all weights from both decoders. Each point represents one neuron. Pearson’s correlation coefficients (*r*) and numbers of neurons (n) are shown. We restricted the comparison of decoder weights across halves to spatially selective neurons, identified using a mutual-information test for position (see Methods).

**Extended Data Fig. 8.**
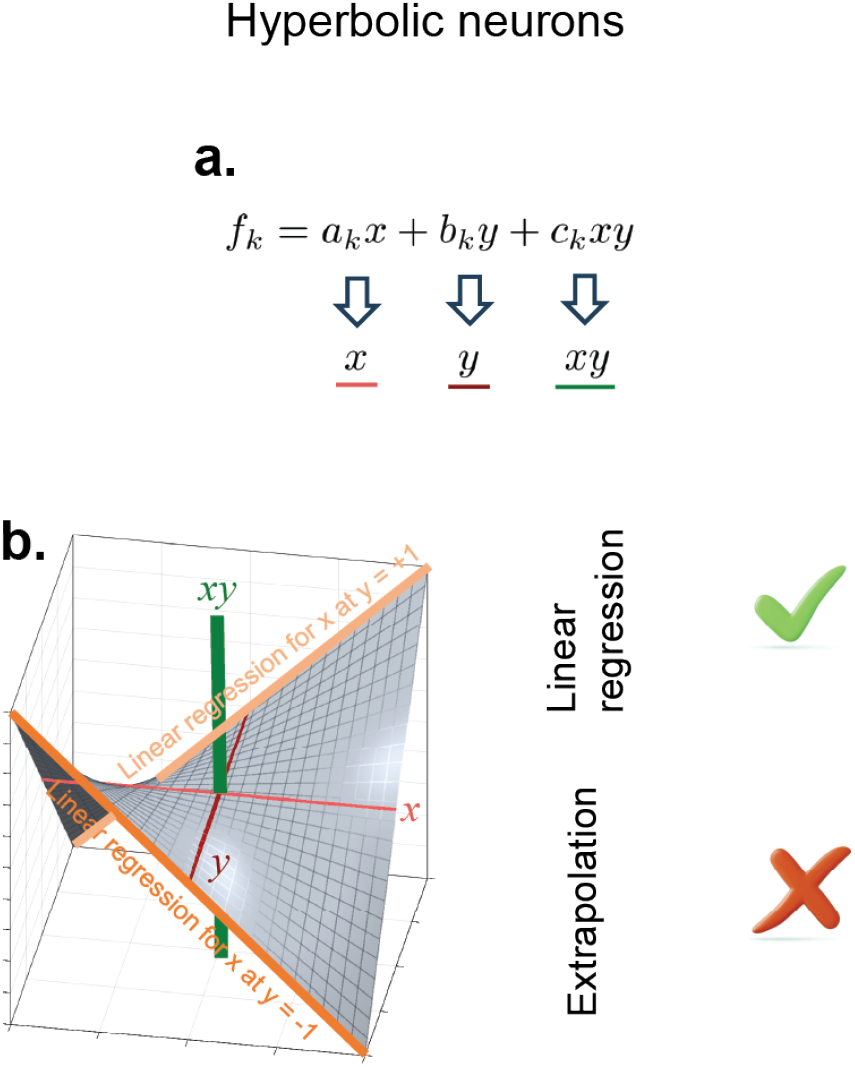
Adding an *xy* term to a linear model prevents linear generalization. **a**, Hyperbolic neurons: the firing rate is a linear function of *x, y*, and *xy*. **b**, The manifold is a hyperbolic paraboloid for which the coding direction of the *x* coordinate (orange lines) varies with *y*, and hence linear regression does not generalize.

**Extended Data Fig. 9.**
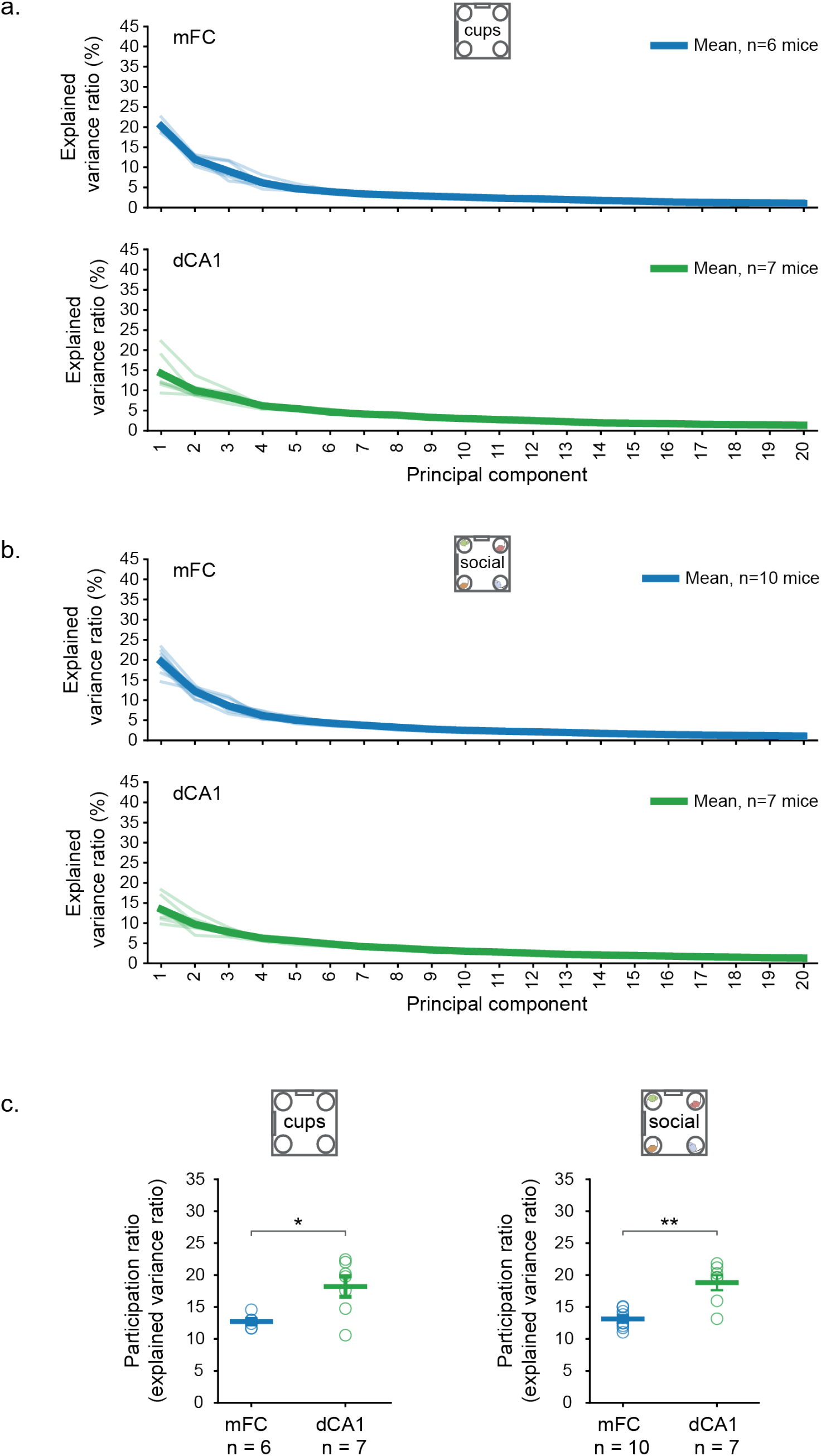
Variance explained by individual principal components. **a-b**. Percentage of variance explained by each of the first 20 principal components derived from Principal Component Analysis of spatial rate maps in the cups assay **(a)** and social assay **(b)**. Thin lines indicate individual mice and thick lines indicate the mean across mice, shown separately for mFC and dCA1. **c**. Participation ratio of the variance explained by each PC in the cups assay (left) and social assay (right). Each circle represents data from one mouse; horizontal lines and error bars indicate the mean ± SEM. Statistical significance across brain areas was assessed using a pairwise Welch’s t-test.

**Extended Data Fig. 10.**
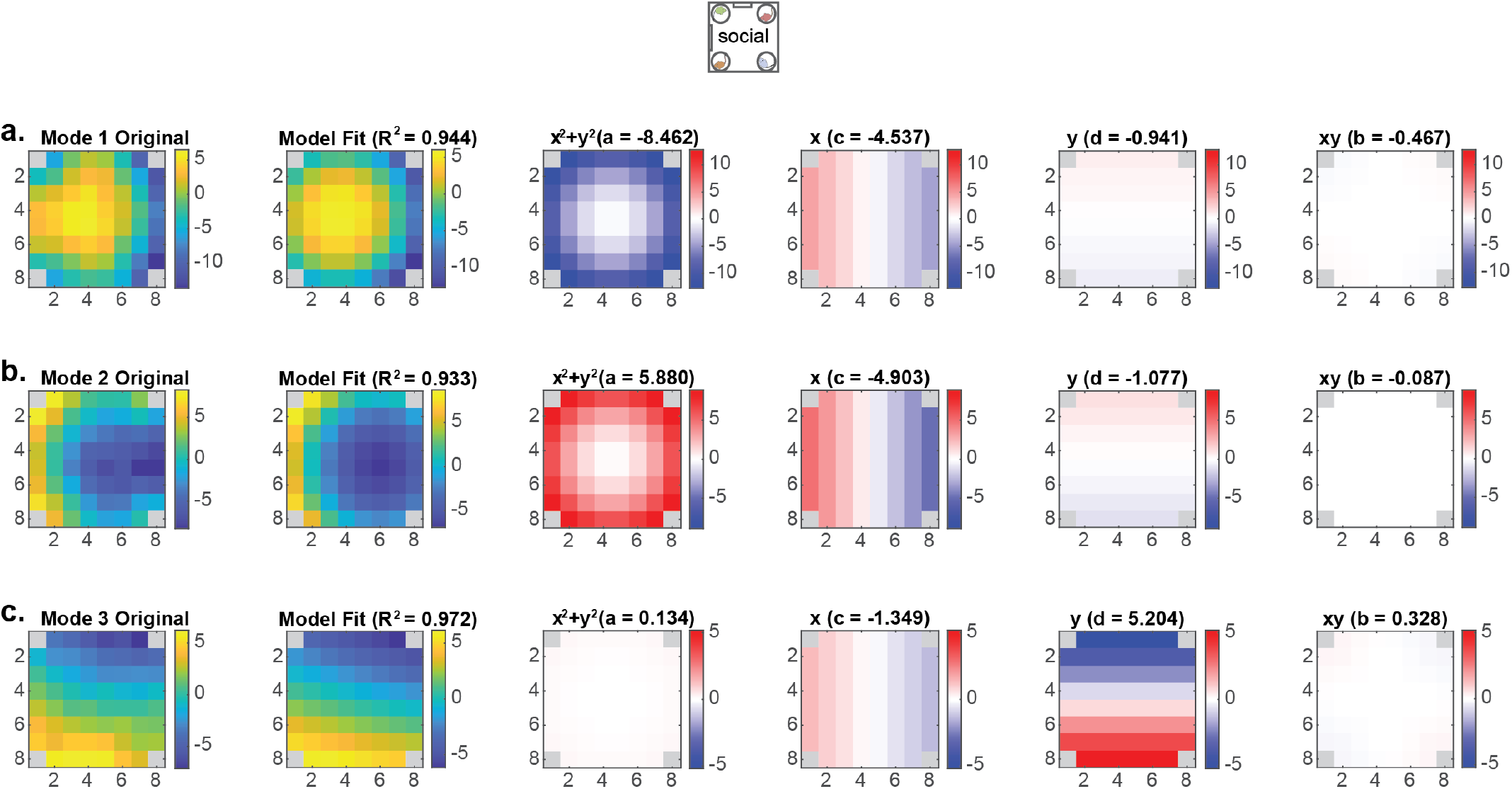
The leading spatial patterns of mFC neural activity in the social assay identified by SVD are explained by a quadratic distance model. **a-c**, The first three spatial modes obtained by singular value decomposition (SVD) of mFC population activity in the social assay.

**Extended Data Fig. 11.**
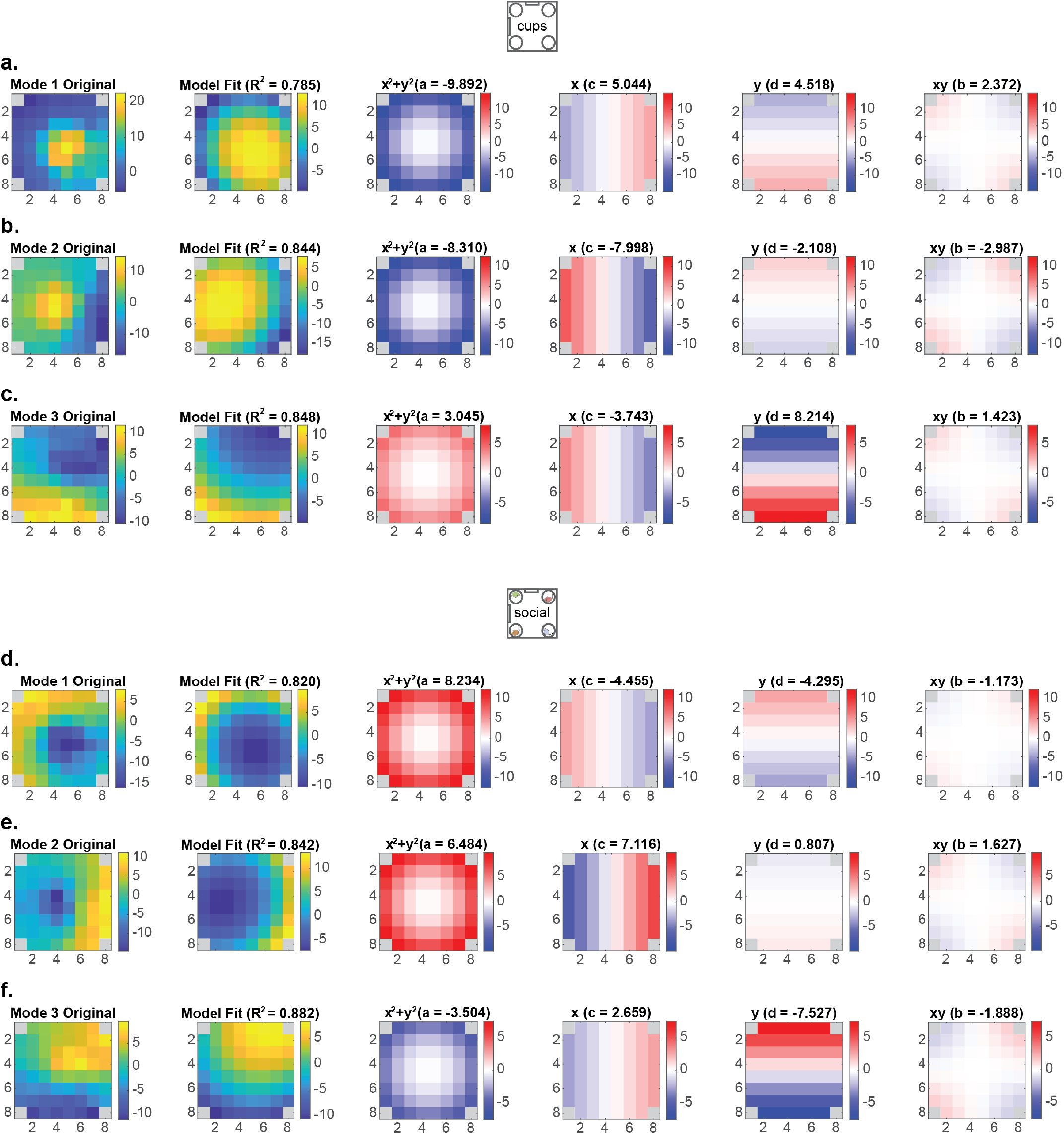
The leading spatial patterns of dCA1 neural activity in the cups and social assays identified by SVD. **a-c**, The first three spatial modes obtained by singular value decomposition (SVD) of dCA1 population activity in the cups assay. **d-f**, Same analysis for the social assay.

**Extended Data Fig. 12.**
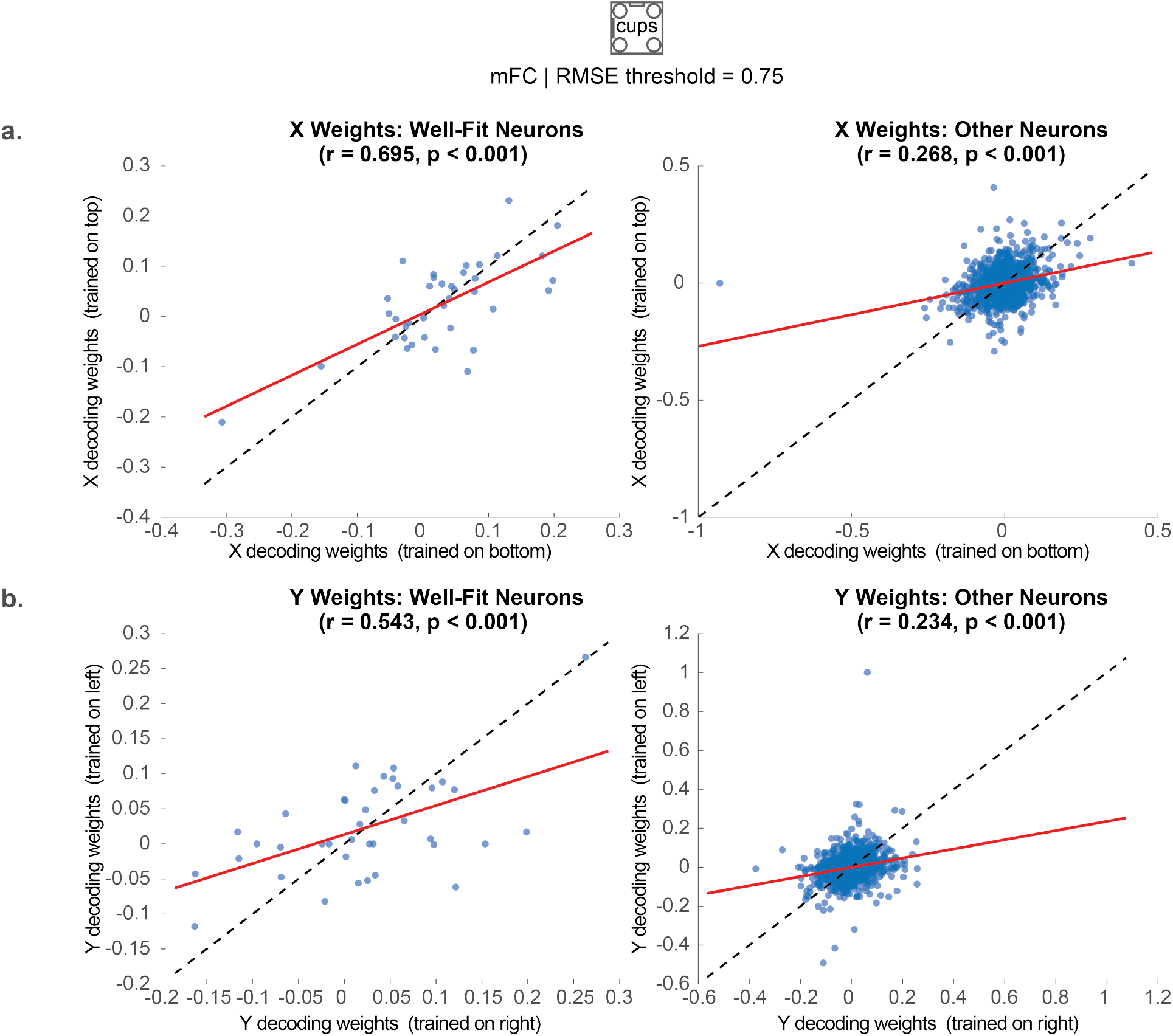
A subset of mFC neurons is well described by a squared distance-based spatial model. **a**, X-position decoder weights obtained from the bottom versus top halves of the arena in the cups assay, shown separately for neurons well fit by the squared distance model (left) and all other neurons (right), using an RMSE threshold of 0.75. **b**, Y-position decoder weights obtained from the right versus left half of the arena in the cups assay, shown separately for neurons well fit by the squared distance model (left) and all other neurons (right), using an RMSE threshold of 0.75. Each point represents one neuron. Red lines indicate linear fits and dashed lines indicate the identity line; Pearson’s *r* and corresponding *p* values are shown above each plot.

**Extended Data Fig. 13.**
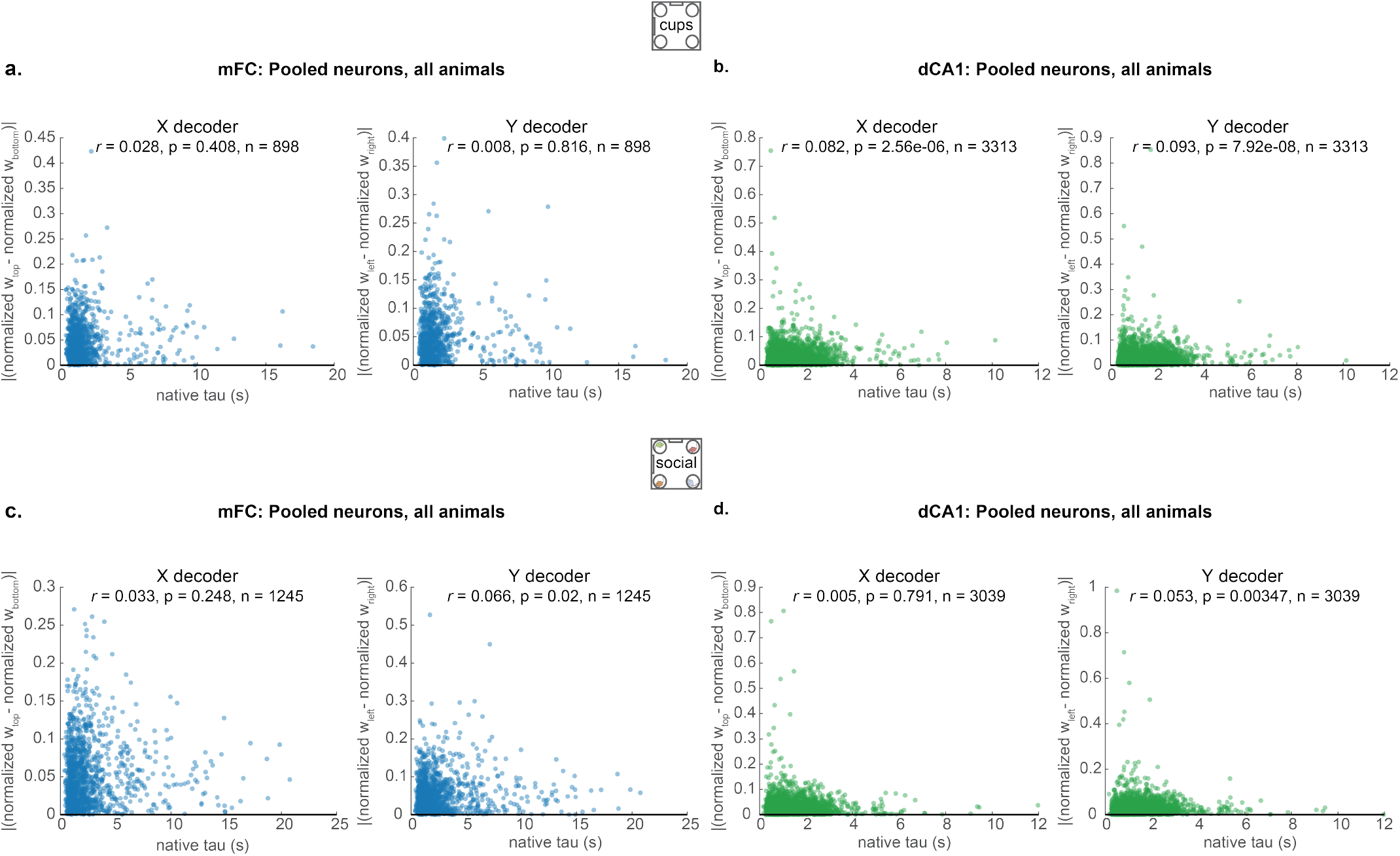
Native calcium decay constant does not strongly predict cross-half decoder-weight consistency. **a**. Relationship between each neuron’s native calcium decay constant (tau) and the absolute difference between its normalized *x*-position decoder weights obtained from decoders trained on the top and bottom halves of the arena (left), or its normalized *y*-position decoder weights obtained from decoders trained on the left and right halves (right), for all mFC neurons pooled across animals in the cups assay. Each point represents one neuron. Pearson’s correlation coefficient (*r*), corresponding *p* value, and number of neurons (*n*) are shown. **b**. Same as **a**, but for the dCA1 neurons in the cups assay. **c**. Same as **a**, but for the mFC neurons in the social assay. **d**. Same as **a**, but for the dCA1 neurons in the social assay.

